# Nicotinamide N-methyltransferase couples inflammation and epigenetic remodeling to hepatic fibrosis

**DOI:** 10.64898/2026.07.30.740978

**Authors:** Sviatlana Sukhanava, Philipp Valina Allo, Yuanyuan He, Lin Chen, Yue Zhu, Sonia Youhanna, Oihane Garcia-Irigoyen, Qi Li, Ewa Ellis, Volker M. Lauschke, Eckardt Treuter, Rongrong Fan

## Abstract

Chronic inflammation drives the progression of metabolic dysfunction-associated steatotic liver disease (MASLD) to fibrosis, yet how inflammatory signals establish profibrotic transcriptional states in hepatocytes remains poorly understood. Here, we identify nicotinamide N-methyltransferase (NNMT) as a previously unrecognized regulator linking inflammatory cytokine signaling to epigenetic and profibrotic responses in hepatocytes. NNMT was selectively induced by IL-1β and IL-6, but not TNFα, and its depletion markedly attenuated cytokine-induced inflammatory and fibrosis-associated transcriptional programs. IL-1β activated NNMT through a JUN/AP-1-dependent pathway, whereas IL-6 sensitized IL-1β-driven responses, an effect partially reversed by NNMT depletion. Integrative transcriptomic and epigenomic analyses revealed that NNMT-dependent gene regulation was associated predominantly with alterations in H3K27 acetylation (H3K27ac) rather than widespread changes in histone methylation. Notably, nicotinamide supplementation closely phenocopied NNMT depletion across cytokine-responsive H3K27ac regions, implicating nicotinamide-dependent regulation in the chromatin response. H3K27ac alterations were strongly concordant with inflammatory and fibrosis-associated transcriptional changes. These findings were further supported in primary human hepatocytes and three-dimensional human liver microtissues, where NNMT depletion reduced cytokine-induced procollagen secretion. Together, our findings identify an inflammation–NNMT–chromatin axis that couples inflammatory cytokine signaling to sustained profibrotic hepatocyte responses and highlight the drug-targetable enzyme NNMT as a potential therapeutic target for limiting MASLD progression and liver fibrosis.

## Introduction

Metabolic dysfunction-associated steatotic liver disease (MASLD) has become the leading cause of liver-related mortality worldwide and is a major contributor to cirrhosis and hepatocellular carcinoma (1). While excessive lipid accumulation initiates disease development, progression from simple steatosis to metabolic dysfunction-associated steatohepatitis (MASH) and liver fibrosis is largely driven by chronic inflammation (2, 3). Although inflammatory responses involve extensive communication between hepatocytes, immune cells and hepatic stellate cells, hepatocytes themselves actively sense inflammatory stimuli and undergo extensive transcriptional reprogramming, producing cytokines, chemokines and profibrotic mediators which amplify hepatic inflammation and activate non-parenchymal cells (2, 4). However, the molecular mechanisms by which inflammatory signaling is translated into persistent pathological transcriptional programs in the hepatocytes remain incompletely understood.

Among the inflammatory cytokines implicated in MASLD progression, interleukin-1β (IL-1β) (5, 6), interleukin-6 (IL-6) (7, 8), and tumor necrosis factor-α (TNFα) (9) are consistently elevated in patients with steatohepatitis and advanced fibrosis. Although IL-1β, IL-6 and TNFα each contribute to hepatic inflammation, they activate distinct intracellular signaling pathways and elicit only partially overlapping transcriptional responses. IL-1β is a potent inducer of inflammatory gene expression through NF-κB and MAPK signaling (5), whereas IL-6 predominantly activates the JAK/STAT3 pathway and modulates both inflammatory and regenerative responses (8). TNFα is another central inflammatory cytokine but displays context-dependent functions in liver injury (9). Increasing evidence suggests that these cytokines are not functionally redundant. Instead, specific cytokine combinations cooperate to amplify inflammatory and fibrogenic responses during chronic liver disease (10, 11). The intracellular mechanisms that integrate these cytokine-specific signals, however, remain poorly understood.

To identify common downstream mediators of inflammatory cytokine signaling, we compared the transcriptional responses of hepatocytes following stimulation with IL-1β, IL-6 and TNFα. Among the differentially regulated genes, nicotinamide N-methyltransferase (*NNMT*) emerged as one of the most robustly induced targets of IL-1β and IL-6, whereas TNFα had little effect on its expression. NNMT is a cytosolic methyltransferase that catalyzes the transfer of a methyl group from S-adenosylmethionine (SAM) to nicotinamide, thereby coupling one-carbon metabolism with nicotinamide adenosine dinucleotide (NAD⁺) metabolism (12, 13). As a result, NNMT elevation may consume NAD⁺ and reduce its endogenous level. Beyond its canonical metabolic function, increasing evidence suggests that NNMT influences cellular epigenetic homeostasis by regulating methyl donor availability and histone methylation (14). Some of the modifications such as H3K27me3 are important regulators defining the chromatin remodeling and gene expression (15). Notably, NAD⁺ serves as an essential cofactor for sirtuin deacetylases and thereby provides a metabolic link to chromatin regulation, whereby alterations in nicotinamide–NAD⁺ metabolism can also influence histone acetylation, including H3K27ac, and consequently transcriptional activity (16, 17). NNMT elevation has been reported in obesity, diabetes, cancer and MASLD, where it has been associated with metabolic dysfunction and disease progression (13, 18–20). Nevertheless, whether NNMT functions as a downstream effector of inflammatory cytokine signaling in hepatocytes remains unknown. Furthermore, it is unclear whether NNMT contributes directly to inflammatory and fibrogenic transcriptional programs through epigenetic remodeling.

In the present study, we identify *NNMT* as a selective downstream target of IL-1β and IL-6 in both human hepatoma cells and primary hepatocytes. We demonstrate that NNMT is required for IL-1β- and IL-6-induced inflammatory and profibrotic gene expression and mediates the ability of IL-6 to potentiate IL-1β-driven inflammatory responses. Furthermore, we show that modulation of *NNMT* expression does not affect histone methylation such as H3K27me3 and H3K4me2 but rather affects histone acetylation markers. H3K27ac is associated with locus-specific epigenetic remodeling at inflammatory genes. The H3K27ac changes induced by nicotinamide treatment or by *NNMT* knockdown are highly correlated, suggesting that the profibrotic effects of NNMT are largely mediated via NAD^+^ deficiency. Together, our findings identify NNMT as a metabolic-epigenetic integrator that couples inflammatory cytokine signaling to chromatin remodeling in hepatocytes, thereby promoting inflammatory and fibrogenic responses relevant to MASLD progression.

## Results

### Inflammatory cytokines promote fibrotic programs via induction of *NNMT* in human hepatic cells and mice

To investigate hepatocyte responses to acute inflammatory stress, HuH-7 cells were stimulated with a cytokine mix consisting of recombinant human IL-1β, TNFα, and IL-6, followed by bulk RNA-seq. Differential expression analysis identified 1,050 significantly upregulated and 1,213 downregulated genes in cytokine mix-treated cells compared with untreated controls (**Figure 1A**). Among the most strongly induced genes were *NNMT*, together with classical inflammatory and fibrosis-associated genes including *CXCL6*, *CXCL8*, *SERPINE1*, *THBS1*, *FRZB*, and *HAS2* (**Figure 1A**). KEGG pathway enrichment analysis confirmed robust activation of inflammatory signaling pathways, including the TNF, NF-κB, IL-17, and cytokine-cytokine receptor interaction pathways (**Figure 1B**). Because NNMT is a well-established metabolic regulator, its induction by inflammatory cytokines may bridge inflammation with metabolic dysregulation in the liver during MASLD progression. To investigate the contribution of NNMT to cytokine-driven transcriptional responses, HuH-7 cells were transfected with control siRNA or siNNMT prior to cytokine mix stimulation and subjected to bulk RNA-seq. Principal component analysis (PCA) revealed that while cytokine treatment shifted the transcriptome along PC1 (48.9% variance), *NNMT* knockdown in the presence of cytokines partially restored the transcriptomic profile (**Figure S1A**) and attenuated the induction of many cytokine-responsive genes, including *IL18*, *IL32*, *SAA1*, *SAA2*, *THBS1*, *PDGFC*, *HAS2*, *CSF1*, *FRZB*, and *CD47* (**Figure 1C**). Comparison of cytokine mix-induced genes with those downregulated following *NNMT* knockdown identified 163 shared genes [SS1](**Figure 1D**). Gene Ontology (GO) analysis of these *NNMT*-dependent genes revealed enrichment of pathways associated with inflammation and liver fibrosis, including TGFβ receptor signaling, extracellular matrix organization, wound healing, inflammatory signaling, cytokine signaling, leukocyte migration, and cell adhesion, all related with development of liver fibrosis (**Figure 1E**).

**Figure 1.**
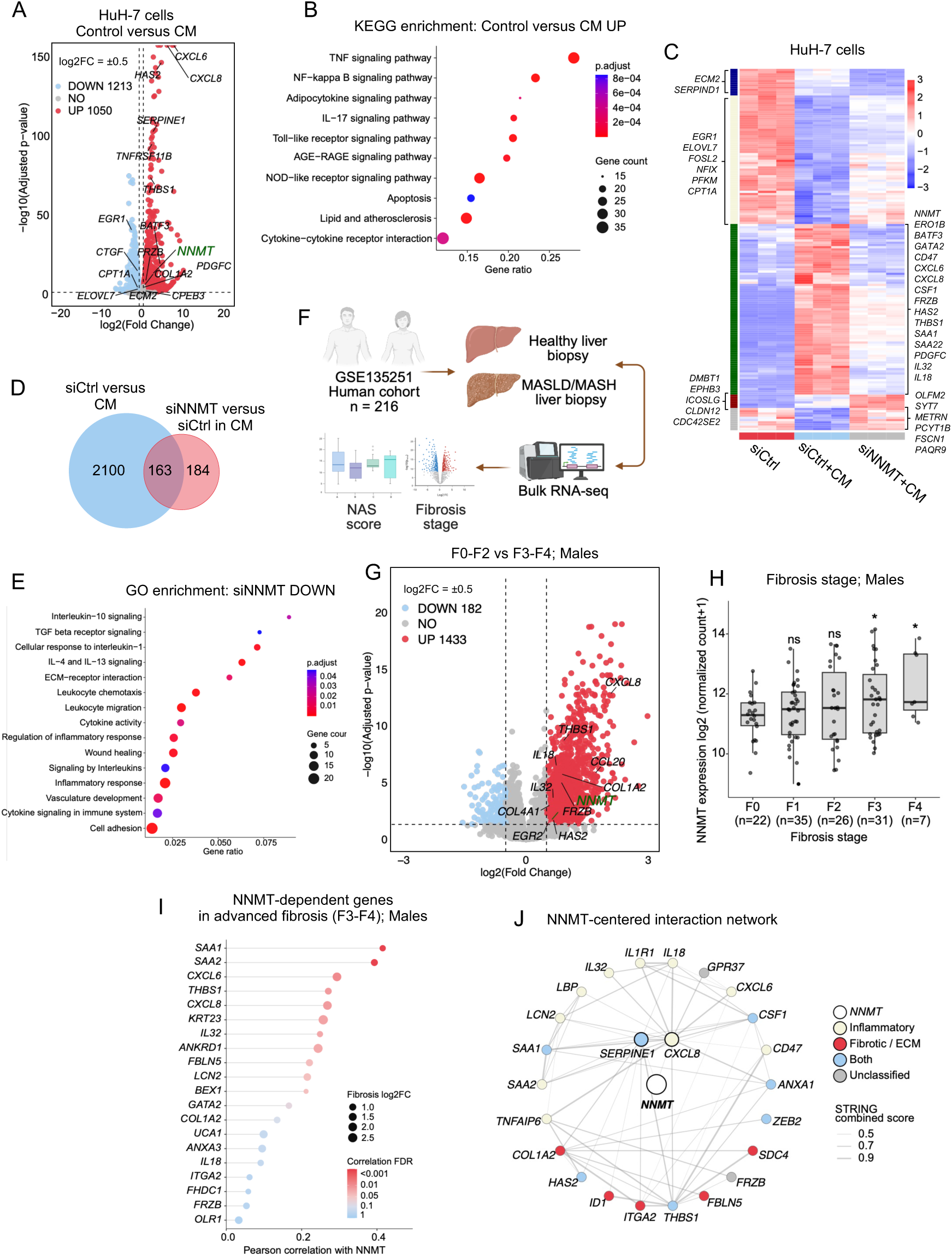
**Pro-inflammatory cytokines induce *NNMT* expression and an inflammatory**, fibrosis-associated transcriptional program in HuH-7 cells. (A) Volcano plot of differentially expressed genes in HuH-7 cells treated with a pro-inflammatory cytokine mix (IL-1β, IL-6 and TNFα; labelled as CM in all main and supplemental figures) versus control. Genes were called at |log2 fold change| > 0.5; red, upregulated (n = 1,050); blue, downregulated (n = 1,213); grey, not significant. Selected inflammatory, fibrosis-associated and metabolic genes are labelled. (B) KEGG enrichment of genes upregulated by CM; dot size indicates gene count and colour indicates adjusted p-value. (C) Heatmap of differentially expressed genes across siCtrl, siCtrl+CM and siNNMT+CM HuH-7 cells; values are row-scaled Z-scores and the left-hand colour bar denotes co-regulated gene groups. Selected genes are labelled. (D) Overlap between CM-induced genes (siCtrl versus CM) and genes reduced by *NNMT* knockdown during CM stimulation (siNNMT versus siCtrl in CM). (E) GO enrichment of genes reduced by *NNMT* knockdown in CM-treated cells; dot size indicates gene count and colour indicates adjusted p-value. (F) Overview of the patient transcriptomic analysis based on the public GSE135251 cohort (n = 216), in which healthy and MASLD/MASH liver biopsies were profiled by bulk RNA-seq and analysed with respect to NAS score and fibrosis stage. Created with BioRender. (G) Volcano plot of genes differentially expressed between early (F0-F2) and advanced (F3-F4) fibrosis in male patients (|log2 fold change| > 0.5; red, upregulated, n = 1,433; blue, downregulated, n = 182); selected inflammatory and fibrosis-associated genes are labelled. (H) *NNMT* expression across fibrosis stages F0-F4 in male patients; each point represents one patient and the number of patients per stage is indicated on the axis. Comparisons are relative to stage F0. (I) Genes correlated with *NNMT* expression in advanced fibrosis (F3-F4) in male patients, ranked by Pearson correlation coefficient; dot size indicates the fibrosis-associated log2 fold change (F3-F4 versus F0-F2) and colour indicates the correlation FDR. (J) NNMT-centred interaction network of inflammatory and fibrotic/ECM genes; node colour denotes functional classification (NNMT, inflammatory, fibrotic/ECM, both, or unclassified) and edge width indicates the STRING combined interaction score. *p < 0.05; ns, not significant.

To evaluate the clinical relevance of these *in vitro* findings, we integrated our experimental data with a public bulk RNA-seq dataset of liver biopsies obtained from 216 individuals spanning healthy controls and patients with MASLD and MASH (21) (**Figure 1F**). We first grouped the patients based on their liver fibrosis stages, with F0-F2 as early and F3-F4 as advanced fibrosis. Differential expression analysis demonstrated that *NNMT*-dependent inflammatory and fibrotic genes identified *in vitro* were also dysregulated in human MASLD and MASH patients. These included *NNMT* itself, as well as multiple *NNMT*-dependent genes (**Figure 1G**). Interestingly, liver *NNMT* mRNA expression increased with liver fibrosis stages in male but not female subjects (**Figure 1H, Figure S1B**). However, when the patients were stratified by NAS score, *NNMT* mRNA levels did not differ significantly among groups in either males or females (**Figure S1C, S1D**). Moreover, we also observed that expression of many NNMT-dependent genes (upregulated by cytokine mix and reduced by *NNMT* knockdown in HuH-7 cells) were also increased in advanced liver fibrosis patients (**Figure 1I**). Based on these integrated analyses, we defined an *NNMT*-centered interaction network using STRING network analysis (22), consisting of extracellular matrix regulators and profibrotic signaling molecules, including *CXCL8*, *SERPINE1*, etc. (Figure 1J).

To determine whether cytokine-induced *Nnmt* expression is conserved *in vivo*, C57BL/6J mice were exposed to lipopolysaccharide (LPS, 1 mg/kg) to induce acute systemic inflammation (**Figure S1E**). Hepatic qPCR analysis confirmed robust induction of *Il1β*, *Il6*, *Tnf*, and *Nnmt* following LPS administration (**Figure S1F**). To further assess the conservation of this transcriptional program, we compared the transcriptional signature identified in cytokine mix-treated HuH-7 cells with a public bulk RNA-seq dataset from livers of LPS-treated mice (23). This analysis revealed substantial overlap between cytokine mix-regulated genes in HuH-7 cells and those altered following LPS treatment in mice, including inflammatory, MASLD and fibrosis-associated genes (**Figure S1G and S1H**). These findings were further supported by an independent public database of liver samples from LPS-treated mice (23) (**Figure S1I**).

Together, these findings demonstrate that inflammatory cytokines induce fibrotic transcriptional signatures in hepatic cells that are partially dependent on NNMT.

### Pro-inflammatory cytokines induce conserved transcriptional programs in both HuH-7 and primary human hepatocytes

To define the contribution of individual pro-inflammatory cytokines to *NNMT* induction and downstream fibrotic transcriptional programs, HuH-7 cells and PHHs from four different donors were stimulated with IL-1β, IL-6, or TNFα separately and analyzed by bulk RNA-seq (**Figure S2A**). PCA demonstrated clear separation of cytokine-treated samples from untreated controls, with IL-1β producing the largest transcriptional shift in both HuH-7 cells and PHHs (**Figure S2B**).

Differential expression analysis of PHHs identified both cytokine-specific and shared transcriptional targets (**Figure 2A**). Interestingly, IL-1β induced the strongest transcriptional response in both PHHs and HuH-7 cells, whereas TNFα regulated relatively small gene subsets (**Figure 2A**), a pattern also observed in HuH-7 cells **(Figure S2C)**. There were also relatively few common genes altered by all the three cytokines, supporting the high cytokine specificity in hepatocytes. Notably, IL-6 induced *NNMT* expression in both PHHs and HuH-7 cells, whereas IL-1β-induced *NNMT* was more robust in HuH-7 cells than in PHHs (**Figure 2A and Figure S2C**). We next compared the NNMT-dependent transcriptional signature in cytokine mix-treated HuH-7 cells with RNA-seq profiles of human PHHs exposed to IL-1β, IL-6, or TNFα. The NNMT-dependent signature showed the strongest overlap with the transcriptional response to IL-1β **(Figure 2B)**. Similar results were also observed in HuH-7 cells (**Figure S2D**). These data suggested that IL-1β is the major cytokine driving the NNMT-associated transcriptional program. KEGG pathway enrichment analysis demonstrated that genes upregulated by IL-1β and TNFα in both cell models were primarily associated with inflammatory pathways, including TNF, NF-κB, chemokine, and IL-17 signaling (**Figure 2C**). In contrast, downregulated genes in both models were primarily associated with metabolic pathways, although fewer significantly enriched pathways were identified (**Figure S2E**). To validate the transcriptomic analyses, representative genes associated with fibrosis, inflammation, and metabolic regulation were analyzed by qPCR in HuH-7 cells (**Figure 2D-F**). Overall, the qPCR data largely recapitulated the RNA-seq findings, further supporting IL-1β as the strongest inducer of these transcriptional programs related to fibrotic and inflammatory responses in the liver cells.

**Figure 2.**
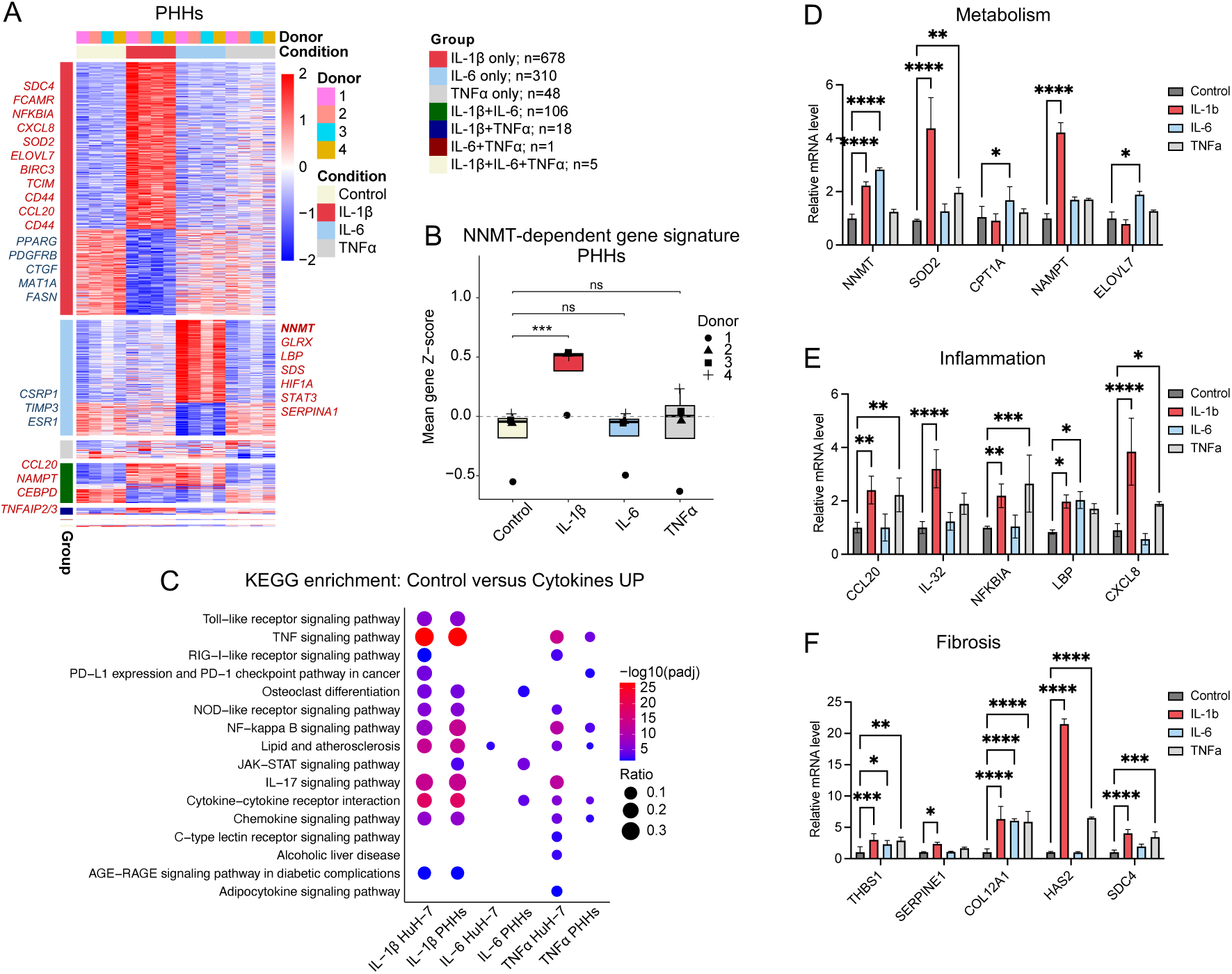
Individual pro-inflammatory cytokines induce distinct but conserved transcriptional programs in primary human hepatocytes. (A) Heatmap of differentially expressed genes in primary human hepatocytes (PHHs) from four donors following IL-1β, IL-6 or TNFα treatment relative to control. Genes are grouped by cytokine-specific and shared responses, with the number of genes per group given in the key; colours indicate row-scaled expression Z-scores. Selected upregulated (red) and downregulated (blue) genes are labelled. (B) *NNMT*-dependent gene signature score in cytokine-treated PHHs, calculated as the mean row-scaled Z-score across the signature genes; each point represents one PHH donor and comparisons are relative to control. (C) KEGG enrichment of genes upregulated by individual cytokines in HuH-7 cells and PHHs; dot size indicates gene ratio and colour indicates -log10 adjusted p-value. (D-F) qPCR validation of representative metabolic (D), inflammatory (E) and fibrosis-associated (F) genes following IL-1β, IL-6 or TNFα treatment in HuH-7 cells. qPCR data are mean ± SD and were analysed by unpaired two-tailed t-test; the signature score in (B) was calculated as the mean Z-score across *NNMT*-dependent genes for each donor and analysed using paired t-tests with Benjamini–Hochberg correction for multiple comparisons. *p < 0.05, **p < 0.01, ***p < 0.001, ****p < 0.0001; ns, not significant.

Comparison of cytokine-responsive genes between PHHs and HuH-7 cells also revealed conserved gene expression changes in response to individual cytokines. IL-6 and TNFα induced very few common genes in HuH-7 and PHHs, while IL-1β triggered much stronger responses, with 169 upregulated and 71 downregulated genes in both HuH-7 and PHHs (**Figure S2F**). Likewise, GO and KEGG analysis of commonly regulated genes identified shared enrichment of inflammatory signaling, cytokine responses, and cell adhesion (**Figure S2G**), demonstrating that the major biological processes activated by inflammatory cytokines are conserved across both hepatocyte models.

Collectively, these findings demonstrate that individual pro-inflammatory cytokines make complementary contributions to the inflammatory response in human hepatocytes. Whereas IL-1β acts as the dominant driver of inflammatory and fibrosis-associated gene expression, *NNMT* expression is controlled mainly by IL-6.

### The transcription factor c-Jun mediates IL-1β-induced activation of *NNMT* and its associated fibrotic regulatory program

To investigate the mechanisms underlying cytokine-induced transcriptional responses at the chromatin level, we first mapped the activated genomic regions upon different cytokine treatment using H3K27ac ChIP-seq and ATAC-seq. The techniques map genome-wide active and highly accessible chromatin regions (24, 25). Following IL-1β, IL-6, and TNFα stimulation in HuH-7 cells, genome-wide analyses identified both shared and cytokine-specific changes in H3K27ac occupancy, including regions that gained or lost acetylation following cytokine treatment (**Figure S3A and S3B**). In contrast, differential chromatin accessibility was comparatively modest, suggesting that cytokine stimulation is accompanied predominantly by remodeling of the active regulatory landscape rather than widespread changes in chromatin accessibility (**Figure S3C**). Representative genome browser tracks illustrated increased H3K27ac enrichment at regulatory regions associated with *NNMT*, *CD47*, *NFKBIA*, *IL18*, and *CCL20*, with IL-1β generally producing the strongest responses, whereas corresponding ATAC-seq profiles showed comparatively limited changes (**Figure 3A**). Similar patterns were observed at additional fibrosis-, inflammatory-, and metabolism-associated loci (**Figure S3D and S3E**).

**Figure 3.**
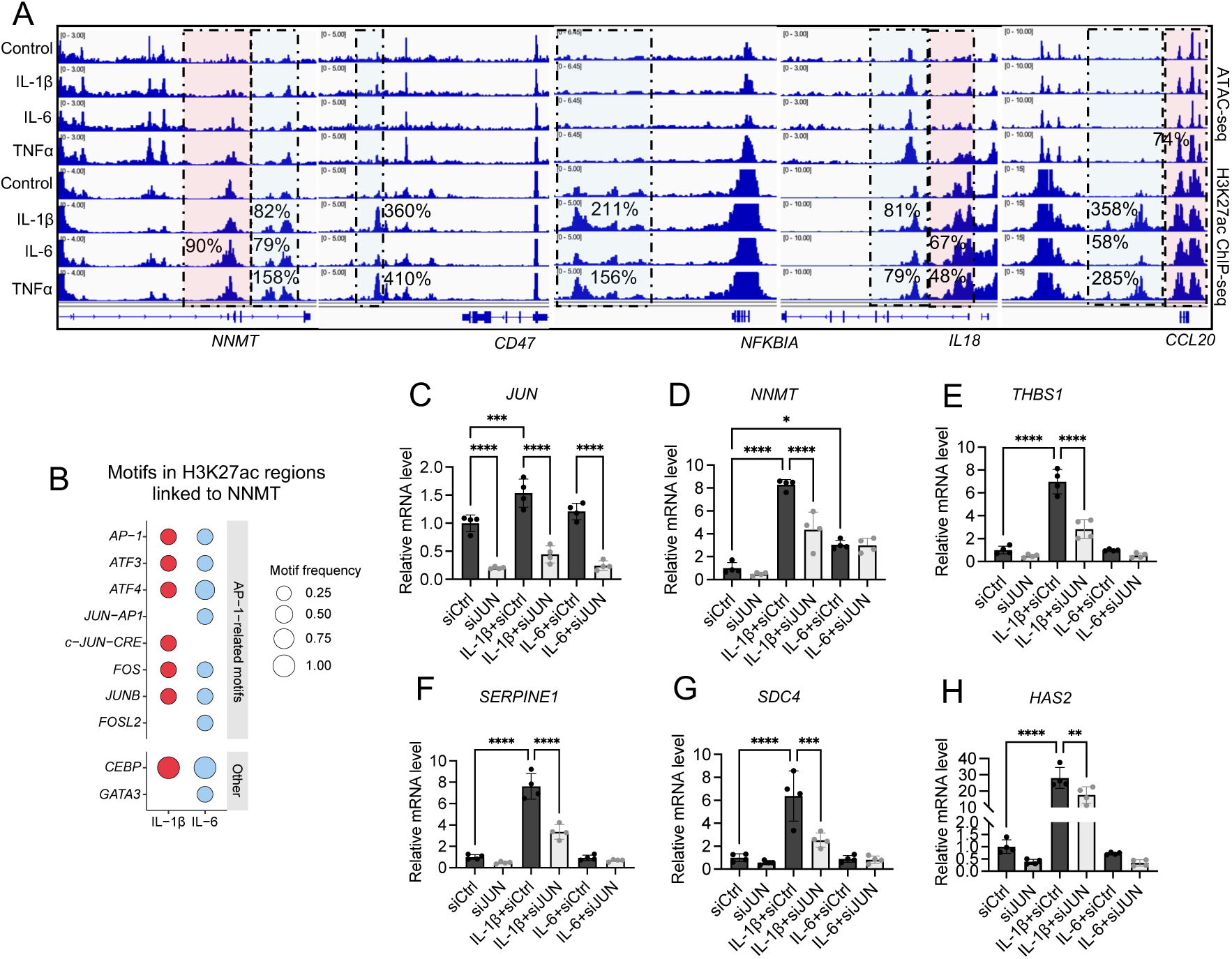
c-Jun mediates IL-1β-induced activation of an NNMT-associated fibrotic regulatory program. (A) Representative genome browser tracks showing ATAC-seq (upper) and H3K27ac ChIP-seq (lower) signal at the *NNMT*, *CD47*, *NFKBIA*, *IL18* and *CCL20* loci following IL-1β, IL-6 or TNFα treatment in HuH-7 cells. Red shading indicates promoter regions and blue shading indicates intronic or intergenic regulatory regions; percentages indicate the change in signal relative to control. (B) Dot plot of transcription factor motifs predicted in H3K27ac regions linked to *NNMT* in IL-1β- and IL-6-treated cells, separated into AP-1-related and other motifs; dot size indicates motif frequency. (C-H) qPCR analysis of *JUN* (C), *NNMT* (D), *THBS1* (E), *SERPINE1* (F), *SDC4* (G) and *HAS2* (H) expression following IL-1β or IL-6 treatment and siJUN-mediated *JUN* knockdown in HuH-7 cells. qPCR data are mean ± SD and were analysed by unpaired two-tailed t-test. *p < 0.05, **p < 0.01, ***p < 0.001, ****p < 0.0001.

To identify transcription factors associated with cytokine-responsive regulation of *NNMT*, we integrated changes in H3K27ac signal with differential gene expression using a manually implemented analytical framework based on IMAGE (26), and subsequently focused on regulatory regions linked to *NNMT*. This analysis identified several AP-1-related motifs under IL-1β and IL-6 stimulation (**Figure 3B**). Among these, the c-Jun-CRE motif was selectively enriched in IL-1β-responsive regions. Consistent with these findings, genome-wide motif enrichment analysis of cytokine-specific H3K27ac-gained regions revealed distinct transcription factor programs across cytokine conditions (**Figure S3 F-H**). IL-1β-responsive regions were characterized by prominent AP-1- and ETS-associated motifs, whereas IL-6-responsive regions showed stronger enrichment of HNF family and nuclear receptor motifs. TNFα-responsive regions displayed comparatively greater enrichment of NF-κB-associated motifs. Together, these findings indicate that individual cytokines engage distinct regulatory programs despite partial overlap in their transcriptional responses. Given that IL-1β emerged as the principal driver of the fibrosis-associated transcriptional program (**Figure 2C and Figure S2E**), the AP-1 family member c-Jun (encoded by the gene *JUN*) was selected for functional validation.

To study the role of c-Jun, HuH-7 cells were transfected with siRNA targeting *JUN* prior to cytokine stimulation. *JUN* knockdown efficiently reduced *JUN* expression under both basal and cytokine-treated conditions (**Figure 3C**). Importantly, *JUN* silencing markedly attenuated IL-1β-induced *NNMT* expression, whereas the induction of *NNMT* by IL-6 was not affected (**Figure 3D**), indicating that IL-1β and IL-6-mediated regulation of *NNMT* occurs through distinct mechanisms.

We next investigated whether c-Jun also regulates downstream fibrosis-associated genes previously identified within the *NNMT*-associated fibrosis module. Silencing *JUN* significantly reduced the IL-1β-mediated induction of *THBS1*, *SERPINE1*, *SDC4*, and *HAS2* (**Figure 3E-H**), whereas the comparatively weaker responses observed following IL-6 stimulation were unaffected. Collectively, these findings identify c-Jun as a critical mediator of IL-1β-induced activation of the *NNMT*-associated fibrotic transcriptional program in hepatocytes.

### IL-6 but not TNFα sensitizes IL-1β-driven transcriptional responses and is partially reversed by *NNMT* knockdown

To determine how individual cytokine contributes to each other’s’ transcriptional regulation, we first compared gene expression following stimulation with each cytokine alone or in combination. Among genes induced by IL-1β alone, 400 were further enhanced upon use of the cytokine mix (50.6% overlap). These genes were classified as sensitization responses (defined as genes which are not induced by IL-6 or TNFα solely but further increased upon combined treatment with IL-1β compared to IL-1β treatment alone) rather than additive effects (genes induced by both cytokines) (**Figure 4A**). In contrast, only a minority of IL-6-(1.8% overlap) or TNFα-responsive genes (6.7% overlap) exhibited additional sensitization upon exposure to the cytokine mix, with most genes showing either additive responses or no further induction (**Figure S4 A and B**). These findings indicate that the transcriptional response to the cytokine mix is characterized predominantly by sensitization of the IL-1β-induced gene program.

**Figure 4.**
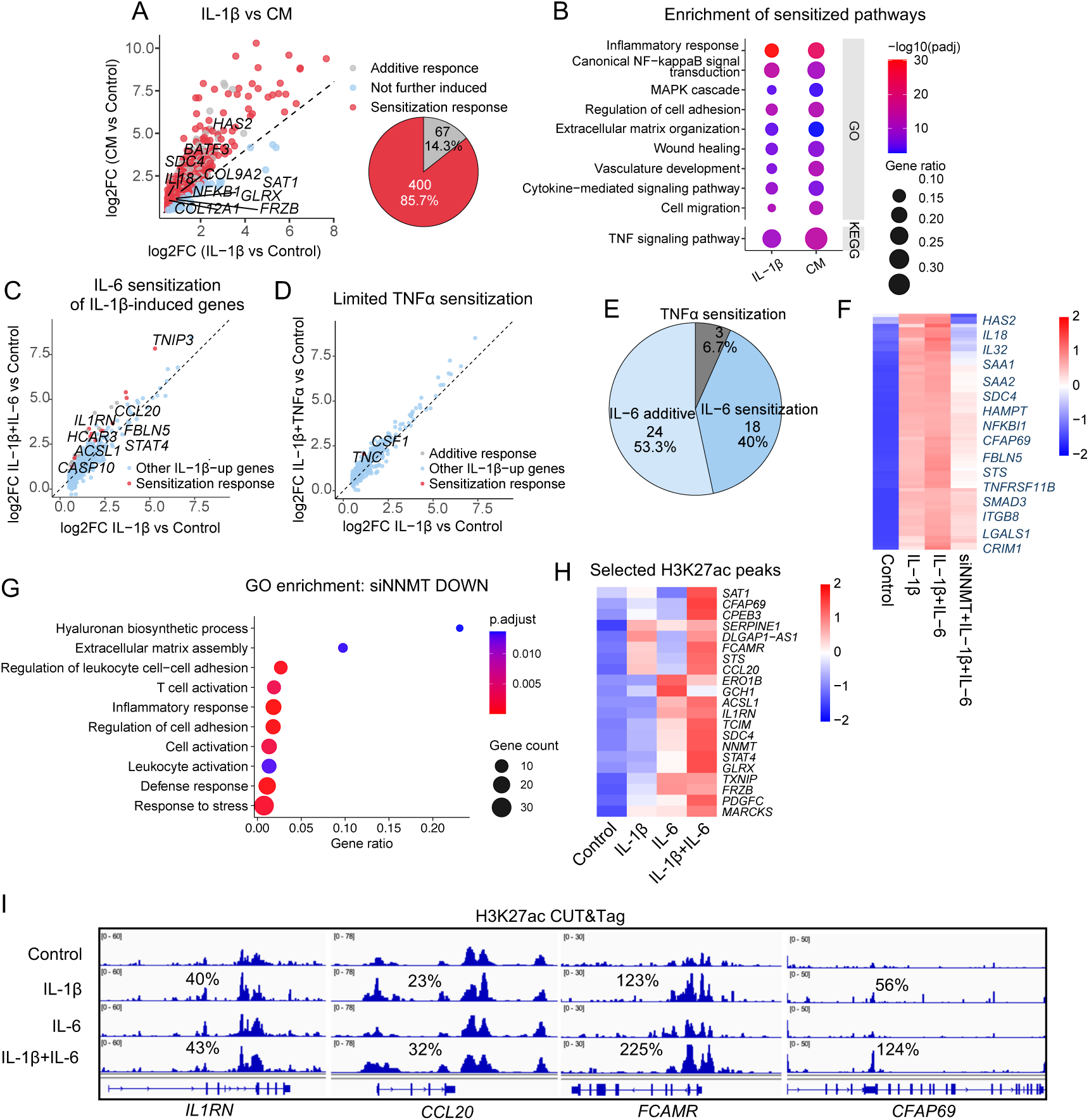
IL-6, but not TNFα, sensitizes IL-1β-driven transcriptional responses, which are partially reversed by *NNMT* knockdown. (A) RNA-seq comparison of IL-1β versus CM responses in HuH-7 cells; genes are classified as additive responses, not further induced, or sensitization responses. The pie chart shows the proportion of sensitized genes (n = 400; 85.7%) among overlapping IL-1β/CM-induced genes. Selected genes are labelled. (B) GO and KEGG enrichment of pathways associated with IL-1β- and CM-sensitized genes; dot size indicates gene ratio and colour indicates -log10 adjusted p-value. (C, D) Scatter plots showing IL-6-mediated sensitization (C) and limited TNFα-mediated sensitization (D) of IL-1β-induced genes in PHHs. (E) Summary of IL-6- and TNFα-modulated responses in PHHs, showing the proportions of IL-6-sensitized (n = 18; 40%), IL-6-additive (n = 24; 53.3%) and TNFα-sensitized (n = 3; 6.7%) genes. (F) Heatmap of genes induced by IL-1β+IL-6 and reduced by *NNMT* knockdown in HuH-7 cells; values are row-scaled Z-scores. (G) GO enrichment of genes reduced by *NNMT* knockdown; dot size indicates gene count and colour indicates adjusted p-value. (H) Heatmap of selected H3K27ac peaks across control, IL-1β, IL-6 and IL-1β+IL-6 conditions from a single PHH donor; values are row-scaled Z-scores. (I) Representative H3K27ac CUT&Tag tracks from a single PHH donor at the *IL1RN*, *CCL20*, *FCAMR* and *CFAP69* loci, showing regulatory regions enhanced by IL-1β+IL-6 relative to IL-1β. Percentages indicate the change in H3K27ac signal relative to control.

Functional enrichment analysis of IL-1β-sensitized genes identified pathways associated with inflammatory responses, NF-κB, MAPK, and cytokine-mediated signaling, wound healing, extracellular matrix organization, and cell migration (**Figure 4B**), indicating that cytokine co-stimulation enhances inflammatory and fibrosis-associated components of the IL-1β transcriptional program.

We next investigated whether a similar sensitization response could be detected in PHHs. Co-treatment with IL-6 selectively enhanced the expression of a subset of IL-1β-induced genes that were not induced by IL-6 alone, consistent with a sensitization response (**Figure 4C**). In contrast, TNFα did not produce a detectable sensitization response and showed only limited additive effects to IL-1β (**Figure 4D**). Although individual gene responses varied between donors, the overall pattern was consistent, with IL-6 accounting for the majority of cooperative effects on IL-1β-induced gene expression, whereas TNFα affected only three genes through additive responses (**Figure 4E**). Together, these findings demonstrate that IL-6, but not TNFα, selectively sensitizes IL-1β-driven transcriptional responses in human hepatocytes.

Given the prominent induction of *NNMT* by inflammatory cytokines, we next investigated whether *NNMT* contributes to maintaining the IL-6-sensitized transcriptional program. RNA-seq analysis in HuH-7 cells identified a subset of genes that were further induced by combined IL-1β and IL-6 stimulation compared with IL-1β alone and whose expression was attenuated following *NNMT* knockdown (**Figure 4F**). Only sensitized genes from both models were selected for the analysis. These genes comprised inflammatory mediators, extracellular matrix-associated genes, and metabolic regulators, indicating that *NNMT* contributes to maintaining both inflammatory and fibrosis-associated transcriptional responses during cytokine co-stimulation. GO analysis further demonstrated enrichment of inflammatory responses, cell adhesion, and extracellular matrix assembly among genes reduced by *NNMT* knockdown (**Figure 4G**), supporting a role for *NNMT* in maintaining inflammatory and profibrotic signaling during cytokine co-stimulation.

Finally, we investigated whether IL-6-mediated transcriptional sensitization was accompanied by changes in chromatin activity. H3K27ac heatmap analysis revealed enhanced acetylation at selected regulatory regions following combined IL-1β and IL-6 stimulation compared with either cytokine alone in PHHs (**Figure 4H**). Representative CUT&Tag profiles from the same donor showed increased H3K27ac enrichment at regulatory regions associated with *IL1RN*, *CCL20*, *FCAMR*, and *CFAP69* following cytokine co-stimulation (**Figure 4I**). Collectively, these findings demonstrate that IL-6 selectively sensitizes IL-1β-driven inflammatory and fibrosis-associated transcriptional programs, with *NNMT* contributing to the maintenance of this amplified transcriptional state.

### NNMT regulates inflammatory cytokine-induced liver fibrosis genes via NAD^+^-dependent epigenetic remodeling in hepatocytes

NNMT elevation consumes both methyl donors and NAD^+^ thereby influencing global histone methylation, acetylation and gene transcription (27). To investigate whether NNMT-dependent metabolic changes are associated with chromatin remodeling during inflammatory stimulation, we first assessed the cellular SAM/SAH ratio following *NNMT* knockdown in the presence or absence of cytokine mix. The SAM/SAH ratio showed no significant difference following *NNMT* knockdown and cytokine mix treatment in the HuH-7 cells (**Figure 5A**).

**Figure 5.**
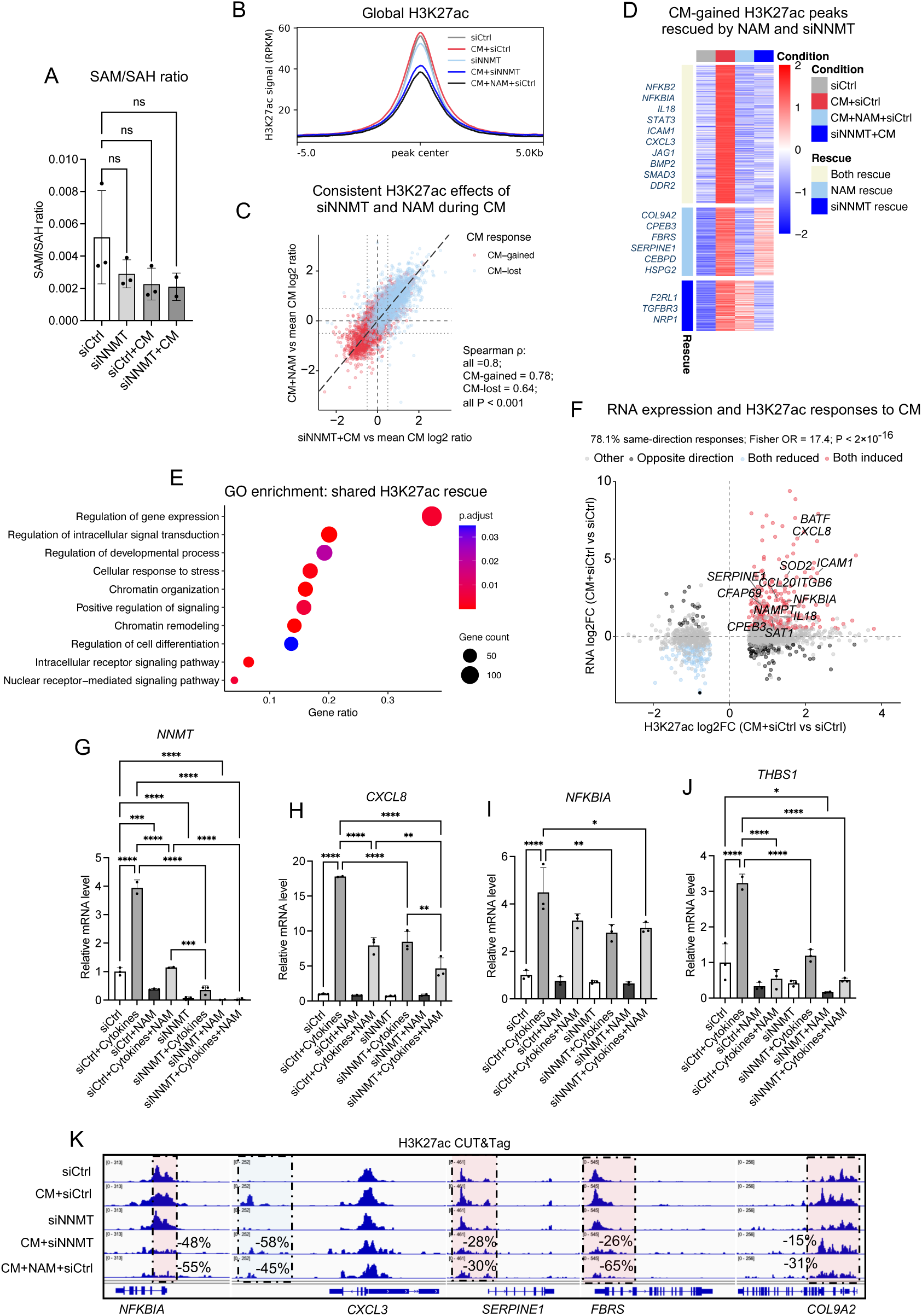
*NNMT* knockdown and NAM are associated with attenuation of cytokine-induced H3K27ac remodeling and inflammatory/fibrotic gene regulation. (A) SAM/SAH ratio in siCtrl, siNNMT, siCtrl+CM and siNNMT+CM HuH-7 cells. Data were analysed by ordinary one-way ANOVA with Dunnett’s multiple-comparison test versus siCtrl. (B) Average H3K27ac signal profile across the global set of analysed H3K27ac peaks in siCtrl, CM+siCtrl, siNNMT, CM+siNNMT and CM+NAM+siCtrl HuH-7 cells. (C) Peak-level comparison of the effects of siNNMT and NAM on CM-responsive H3K27ac regions. Values represent log2 ratios of CM+siNNMT or CM+NAM+siCtrl relative to mean CM+siCtrl signal; CM-gained and CM-lost peaks are indicated separately. Spearman correlations are shown (all peaks, ρ = 0.80; CM-gained, ρ = 0.78; CM-lost, ρ = 0.64; all p < 0.001). (D) Heatmap of CM-gained H3K27ac peaks that were reduced by NAM and/or siNNMT; the right-hand annotation indicates whether rescue was observed with NAM, siNNMT or both. Rescue was defined descriptively as a log2 ratio ≤ −0.5 relative to mean CM+siCtrl signal. Selected inflammatory and fibrosis/remodeling-associated genes are indicated. (E) GO enrichment of genes associated with CM-gained H3K27ac peaks jointly reduced by both siNNMT and NAM (shared H3K27ac rescue). (F) Integration of CM-responsive RNA-seq and H3K27ac CUT&Tag data. Directional concordance was 78.1% of same-direction responses (Fisher’s exact test, odds ratio = 17.4; p < 2 × 10^-^¹⁶). Selected inflammatory, metabolic and fibrosis-associated genes are labelled. (G-J) qPCR analysis of *NNMT* (G), *CXCL8* (H), *NFKBIA* (I) and *THBS1* (J) expression following cytokine stimulation, *NNMT* knockdown and NAM treatment in HuH-7 cells. (K) Representative H3K27ac CUT&Tag genome-browser tracks at the *NFKBIA*, *CXCL3*, *SERPINE1*, *FBRS* and *COL9A2* loci. For H3K27ac CUT&Tag, biological replicate numbers are siCtrl, n = 3; CM+siCtrl, n = 3; siNNMT, n = 2; CM+siNNMT, n = 1; and CM+NAM+siCtrl, n = 1. Percentages indicate the change in H3K27ac signal in CM+siNNMT and CM+NAM+siCtrl relative to mean CM+siCtrl signal. qPCR data are presented as mean ± SD. Statistical comparisons were performed using one-way ANOVA with multiple-comparison correction. *p < 0.05, **p < 0.01, ***p < 0.001, ****p < 0.0001; ns, not significant.

Because the SAM/SAH ratio reflects cellular methyl-donor levels, we next examined whether modulation of NNMT was accompanied by changes in histone methylation levels. In an *NNMT* overexpression experiment, western blotting showed largely comparable global levels of the examined histone methylation marks across conditions (**Figure S5A**). Similarly, genome-wide CUT&Tag profiles showed largely overlapping average H3K4me2 and H3K27me3 signal distributions across experimental conditions (**Figure S5 B and C**), indicating that NNMT-mediated transcriptional regulation did not rely on its role of methyl donor consumption. Locus-specific alterations were also checked. H3K27me3 changes were restricted to a relatively small number of regions, with representative changes detected at inflammatory and metabolic loci such as *CXCL12*, *TAC1* and *LDB2* (**Figure S5D**). H3K4me2 showed more extensive locus-specific remodeling, including altered signal at fibrosis- and cytokine-response-associated loci such as *PDGFC*, *FRZB*, *TNFRSF11B*, *STAT4* and *CRIM1* (**SI Appendix Figure S5E**). These epigenetic alterations were either inconsistent with gene expression changes or with the hypothesis of reduced methyl levels upon *NNMT* overexpression. Based on these data, we conclude that NNMT-associated effects on histone methylation were not involved in its transcriptional regulation of liver fibrosis genes.

Because NNMT indirectly affects the NAD^+^ level by consuming endogenous nicotinamide, and that NAD^+^ is a major activator of deacetylases such as sirtuins, which may act on histone acetylation levels (16). We next checked whether *NNMT* knockdown was associated with changes in H3K27ac, a mark more closely linked to active regulatory regions and cytokine-responsive transcription (28). Average H3K27ac profiles across the analyzed peak set showed an increased mean signal following cytokine mix stimulation versus control, and lower signal in the cytokine mix+siNNMT and cytokine mix+NAM conditions (**Figure 5B**). The global changes in H3K27ac support the hypothesis that NNMT-dependent regulation of liver fibrosis genes is linked to alterations in endogenous NAD+ levels. To prove this, we compared the H3K27ac changes in both *NNMT* knockdown and NAM supplemented HuH-7 cells. Strikingly, the responses to siNNMT and NAM showed a strong positive correlation across all cytokine mix-responsive peaks genome-wide (Spearman ρ = 0.80), with similar relationships among cytokine mix-gained (ρ = 0.78) and cytokine mix-lost (ρ = 0.64) regions (**Figure 5C**). Thus, despite representing distinct interventions, *NNMT* knockdown and NAM produced highly overlapping changes across the cytokine mix-responsive H3K27ac landscape.

We next focused on cytokine mix-gained H3K27ac regions and identified peaks at which the cytokine mix-induced increase was opposed by siNNMT, NAM, or both interventions. A substantial subset of these regions was attenuated by both treatments, while additional peaks showed preferential responses to either NAM or *NNMT* knockdown (**Figure 5D**). These regions were associated with inflammatory signaling genes, including *NFKB2*, *NFKBIA*, *IL18*, *STAT3*, *ICAM1* and *CXCL3*, as well as genes implicated in extracellular matrix regulation and tissue remodeling, including *JAG1*, *BMP2*, *SMAD3*, *DDR2*, *SERPINE1*, *HSPG2*, *TGFBR3* and *NRP1*. Additional representative H3K27ac profiles at *NFKB2*, *STAT3*, *CEBPD* and *SMAD3* further illustrated locus-specific attenuation of cytokine mix-induced acetylation following *NNMT* knockdown and NAM treatment (**Figure S5F**).

GO enrichment analysis of genes associated with cytokine mix-gained regions that were jointly reduced by siNNMT and NAM revealed enrichment of inflammatory pathways including regulation of gene expression, intracellular signal transduction, cellular stress responses, developmental and differentiation programs, and chromatin organization and remodeling (**Figure 5E**). These results indicate that the shared H3K27ac response to siNNMT and NAM encompasses a broad regulatory component rather than a single narrowly defined pathway.

To determine whether cytokine mix-induced H3K27ac changes were associated with transcriptional output, we integrated cytokine mix-responsive CUT&Tag regions with RNA-seq data obtained under the corresponding cytokine condition. Genes significantly altered in both datasets showed pronounced directional concordance, with 78.1% exhibiting changes in the same direction (**Figure 5F**). Concordantly induced genes included inflammatory and stress-response regulators such as *BATF*, *CXCL8*, *ICAM1*, *NFKBIA* and *IL18*, together with fibrosis- and remodeling-associated genes including *SERPINE1* and *ITGB6*. These findings indicate that cytokine mix-dependent H3K27ac remodeling is closely associated with the transcriptional response to inflammatory stimulation.

We next examined selected NNMT-, inflammatory-, and fibrosis-associated transcripts following *NNMT* knockdown and NAM supplementation. Cytokine mix strongly induced *NNMT*, *CXCL8*, *NFKBIA* and *THBS1* expression, whereas *NNMT* knockdown and NAM attenuated selected cytokine mix-induced responses to varying degrees (**Figure 5G-J**). Representative H3K27ac CUT&Tag profiles further demonstrated locus-specific attenuation of cytokine mix-induced acetylation at inflammatory and remodeling-associated regions, including *NFKBIA*, *CXCL3* and *SERPINE1*, together with *FBRS* and *COL9A2* (**Figure 5K**).

Collectively, these findings highlight distinct chromatin responses across the two experimental settings. In the *NNMT* overexpression model, average H3K4me2 and H3K27me3 profiles remained largely similar across conditions, with comparatively modest locus-specific changes. In contrast, H3K27ac profiling following *NNMT* knockdown revealed substantially greater condition-dependent differences during cytokine mix stimulation, and these changes were highly overlapping with those observed following NAM treatment. The alignment between cytokine mix-responsive H3K27ac changes and transcriptional responses further supports an association between NNMT-dependent metabolic regulation, active chromatin states, and inflammatory and fibrosis-associated gene expression.

### *NNMT* knockdown attenuates fibrogenic responses in human liver microtissue

To evaluate the functional relevance of NNMT in human liver models, we investigated its role in patient-derived primary human liver microtissues (**Figure 6A**). The liver microtissues were transfected with control or *NNMT*-targeting siRNA and stimulated with IL-1β, IL-6, TNFα, or a cytokine mix before assessment of intracellular lipid accumulation and procollagen secretion. MASH liver microtissues showed a significant increase in intracellular triglyceride and collagen levels as previously reported (29). Surprisingly, *NNMT* knockdown had only limited effects on intracellular lipid accumulation, although cytokine-dependent differences were observed (**Figure 6B and Figure S6 A-E**). In contrast, procollagen secretion was markedly increased by IL-1β stimulation and the cytokine mix, but not by IL-6 and TNFα (**Figure 6C**). Importantly, *NNMT* knockdown significantly reduced procollagen secretion in IL-1β- and cytokine mix-treated liver microtissues, demonstrating attenuation of IL-1β-driven profibrogenic responses.

**Figure 6.**
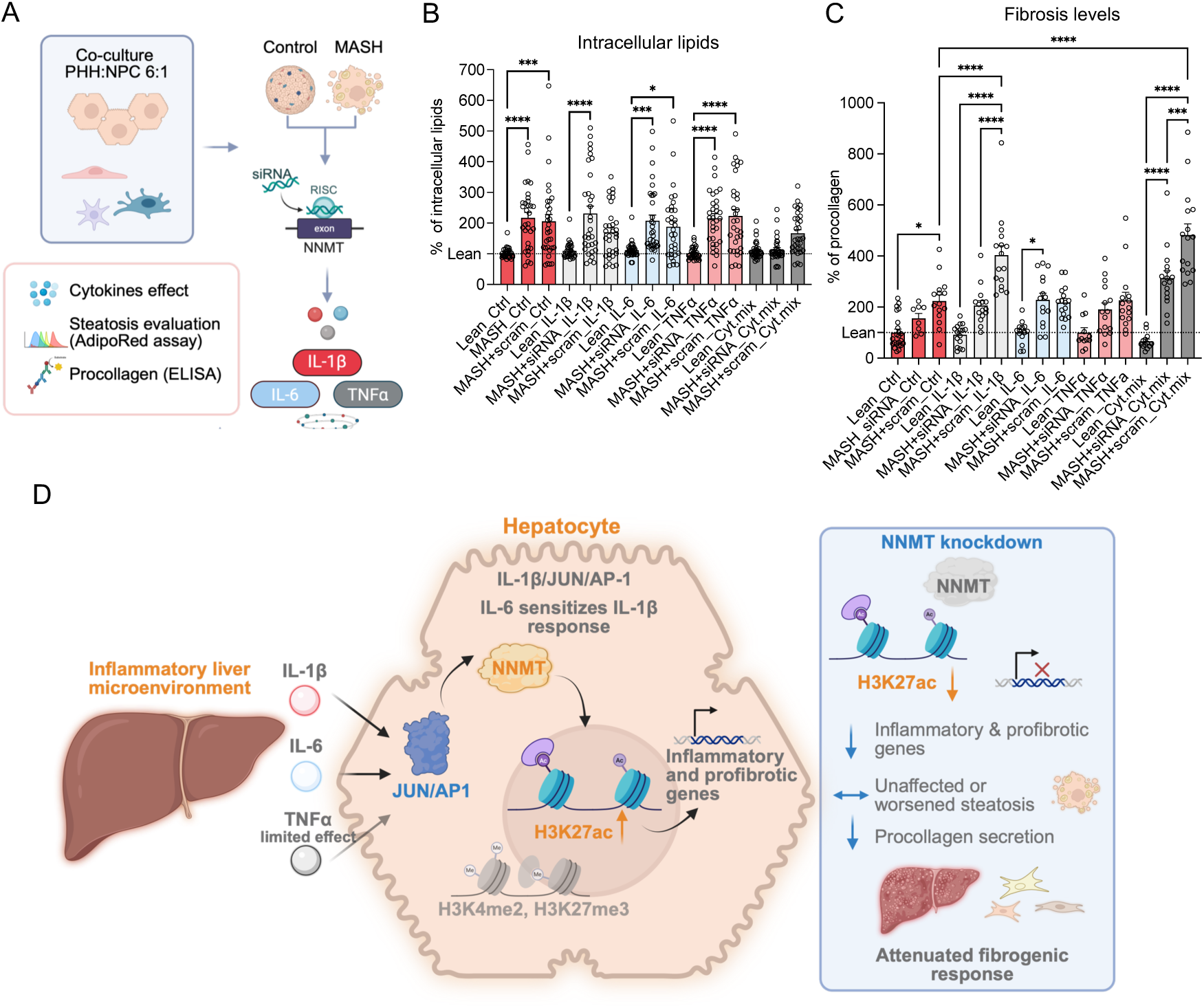
*NNMT* knockdown attenuates fibrogenic responses in human liver spheroids. (A) Schematic overview of the co-culture liver spheroid model containing PHHs and non-parenchymal cells (NPCs) at a 6:1 ratio, with steatosis induction (MASH conditions), cytokine stimulation, siRNA-mediated NNMT knockdown, and AdipoRed and procollagen ELISA readouts. Created with BioRender. (B) Quantification of intracellular lipid accumulation by AdipoRed assay in liver spheroids under lean and MASH conditions with scrambled control siRNA (scram) or *NNMT* siRNA, in the absence or presence of IL-1β, IL-6, TNFα or CM; values are normalised to lean spheroids. (C) Quantification of fibrosis-associated procollagen levels by ELISA in liver spheroids under the same treatment conditions; values are normalised to lean spheroids. (D) Proposed model summarising cytokine- and *NNMT*-associated regulation of hepatocyte inflammatory and fibrogenic responses. IL-1β/JUN/AP-1 signaling promotes inflammatory and profibrotic gene expression through increased H3K27ac, while IL-6 sensitizes the IL-1β response and contributes to *NNMT* induction and TNFα has limited effects. *NNMT* knockdown attenuates inflammatory and profibrotic gene expression and procollagen secretion, whereas steatosis is unaffected or worsened. Created with BioRender. Data are presented as mean ± SD. Statistical comparisons were performed using one-way ANOVA with multiple-comparison correction. *p < 0.05, **p < 0.01, ***p < 0.001, ****p < 0.0001.

Together, these findings identify NNMT as a factor contributing to cytokine-driven amplification of fibrogenic responses in human liver spheroids and further support its association with fibrosis-related processes in human MASLD. Based on the combined transcriptional, epigenetic, and functional data, we propose a model in which inflammatory cytokine signaling, particularly IL-1β through JUN/AP-1 and IL-6-dependent sensitization, promotes hepatocyte *NNMT* expression and is accompanied by cytokine- and NNMT-associated chromatin remodeling. *NNMT* knockdown attenuates inflammatory and profibrotic gene expression, and procollagen secretion, resulting in an overall reduction of the fibrogenic phenotype (**Figure 6D**).

## Discussion

Chronic inflammation is a major determinant of disease progression from steatosis to MASH and liver fibrosis (3). Inflammatory cytokines have long been recognized as key drivers of hepatocyte dysfunction. However, the intracellular mechanisms that integrate cytokine signaling into sustained pathological transcriptional responses in MASLD remain incompletely understood. In the present study, we identify NNMT as a previously unrecognized downstream target selectively induced by IL-1β or IL-6, but not TNFα, in HuH-7 cells and PHHs. We further demonstrate that NNMT is required for inflammatory and profibrotic transcriptional programs induced by IL-1β and IL-6 and mediates the ability of IL-6 to sensitize IL-1β-dependent inflammatory responses. Finally, we provide evidence that NNMT regulates histone acetylation but not methylation at specific inflammatory gene loci, suggesting that NNMT couples inflammatory cytokine signaling to epigenetic remodeling in hepatocytes.

A notable finding of this study is the differential regulation of NNMT by IL-1β, IL-6 and TNFα in hepatocytes. While IL-6 robustly induced NNMT expression in both HuH-7 cells and PHHs, IL-1β increased NNMT only in HuH-7 cells, whereas TNFα failed to induce NNMT and elicited only limited transcriptional responses. This difference likely reflects distinct signaling mechanisms and the requirement for sustained or cooperative transcriptional activation in hepatocytes (30). HuH-7 cells may possess a more permissive chromatin landscape that facilitates IL-1β-responsive gene expression, whereas IL-6 activates the conserved JAK-STAT3 pathway (31), which directly regulates NNMT transcription. Consistent with this, JUN knockdown selectively attenuated IL-1β-induced, but not IL-6-induced, *NNMT* expression, suggesting that IL-1β and IL-6 converge on NNMT through distinct upstream transcription factors. In contrast, TNFα primarily activates transient NF-κB signaling that is rapidly restrained by negative feedback regulators (32) and often requires cooperation with other cytokines to drive robust inflammatory gene expression in hepatocytes (10).

Hepatocytes are exposed to multiple inflammatory cytokines during chronic liver injury, and previous studies have shown that IL-1β promotes NF-κB-dependent chromatin remodeling, thereby facilitating STAT3 recruitment and sensitizing hepatocytes to IL-6 signaling (10). Previous work in liver spheroids showed that IL-1β exposure resulted in sustained hepatocyte dedifferentiation and NF-κB activation by IL-1β or TNFα was required for the stimulation of human hepatocyte regeneration while IL-6 only amplified cell cycle reentry (33, 34). Consistent with this model, we found that IL-6 alone regulated relatively fewer genes but markedly potentiated IL-1β-induced inflammatory gene expression, with approximately 50% of IL-1β-responsive genes showing further induction following co-stimulation. Importantly, *NNMT* knockdown substantially attenuated this transcriptional sensitization, identifying NNMT as a non-canonical downstream mediator of IL-1β-IL-6 crosstalk that amplifies inflammatory and profibrotic transcriptional programs during chronic liver injury.

NNMT has traditionally been studied as a metabolic enzyme that catalyzes the methylation of nicotinamide using S-adenosylmethionine (SAM) as a methyl donor, thereby linking one-carbon metabolism with NAD⁺ homeostasis (19, 35). NNMT is highly expressed in the liver, but its physiological role in hepatic metabolism has remained controversial (19). Previous studies suggest that the contribution of liver NNMT to basal hepatic methyl-donor homeostasis is relatively limited because GNMT is the dominant hepatic methyltransferase (35). In contrast, increasing evidence indicates that NNMT is functionally important during pathological conditions, including MASLD, alcohol-associated liver disease and hepatocellular carcinoma (12, 13, 18, 19). Our findings provide a potential explanation for this context-dependent role by identifying inflammatory cytokines as upstream regulators of *NNMT* expression. Previous studies have demonstrated that hepatic inhibition or deletion of NNMT also protects against MASLD by reducing hepatic steatosis, inflammation and fibrosis, supporting a role for NNMT in metabolic homeostasis and disease progression (13, 18, 35). In contrast, *NNMT* knockdown in our human liver spheroid model selectively reduced collagen production with relatively minor effects on oleate/palmitate-induced steatosis. Rather than representing a discrepancy, these findings likely reflect distinct stages and biological contexts of disease. Previous *in vivo* studies primarily investigated the contribution of NNMT to the initiation and progression of metabolic dysfunction, where long-term modulation of NNMT influences hepatic lipid metabolism, NAD⁺ homeostasis and systemic insulin sensitivity (13, 18, 20, 35). In contrast, our liver spheroid model represents a lipotoxic environment in which steatosis is driven by sustained oleate and palmitate overload(29). Under these conditions, hepatocyte lipid accumulation may occur largely independently of NNMT, whereas inflammatory and fibrogenic responses remain dependent on NNMT-mediated transcriptional reprogramming. Consistent with this interpretation, *NNMT* knockdown preferentially suppressed fibrosis markers while having minimal effects on lipid accumulation. Together, these findings suggest that NNMT may play distinct roles during disease progression: contributing to metabolic dysfunction during the early stages of MASLD while functioning as a cytokine-inducible metabolic-epigenetic regulator that amplifies inflammatory and fibrogenic responses once steatosis has been established.

Recent studies have also shown that NNMT is a regulator of cellular epigenetic states through its effects on methyl donor availability. In cancer and metabolic diseases, increased NNMT activity has been proposed to create a ‘methylation sink’, leading to widespread alterations in histone methylation and transcriptional regulation. Interestingly, *NNMT* overexpression did not alter common histone methylation markers such as H3K27me3 and H3K4me2 but rather changed H3K27ac in human liver cells. These findings suggest that the methyl donor consumption may not contribute to the NNMT-induced liver fibrotic remodeling. Previous studies have also reported that NNMT regulates NAD⁺ homeostasis through nicotinamide methylation (13, 36). Increased NNMT activity has been linked to enhanced sirtuin deacetylase activity by reducing intracellular nicotinamide, a physiological inhibitor of sirtuin deacetylases such as SIRT1 (37). SIRT1 has been implicated in chromatin remodeling through histone deacetylation (38). Future studies investigating H3K27ac dynamics and SIRT1 activity will be important to define the epigenetic mechanisms linking NNMT to inflammatory gene activation.

Our study has several strengths. We demonstrate selective induction of NNMT in both hepatoma cells and primary hepatocytes, combine transcriptomic and epigenetic analyses with functional knockdown experiments, and consistently identify NNMT as a critical mediator of inflammatory responses induced by IL-1β and IL-6. Importantly, our study primarily employed human-relevant experimental systems, including primary human hepatocytes and 3D human liver microtissues, enabling the role of NNMT to be examined in a physiologically relevant human context. While previous studies using genetic mouse models have reported improved metabolic phenotypes following hepatic NNMT depletion, NNMT knockdown in our human liver microtissues did not substantially improve metabolic parameters but instead markedly reduced fibrotic responses, consistent with the inflammatory and profibrotic transcriptional programs identified by our transcriptomic analyses. These findings suggest that, in human liver, NNMT may have a particularly important role in regulating inflammatory and fibrotic responses rather than primarily controlling metabolic homeostasis. Nevertheless, several limitations should be acknowledged. Although our epigenetic analyses demonstrate significant changes in histone acetylation and chromatin occupancy, the direct biochemical mechanisms linking NNMT activity to chromatin remodeling remain incompletely defined.

In conclusion, our findings identify NNMT as a cytokine-inducible metabolic-epigenetic integrator that selectively couples IL-1β and IL-6 signaling to chromatin remodeling and inflammatory transcriptional reprogramming in hepatocytes. Rather than functioning solely as a metabolic enzyme, NNMT appears to translate extracellular inflammatory cues into sustained epigenetic and transcriptional changes that promote inflammatory and profibrotic responses. These findings provide new mechanistic insight into how chronic inflammation is maintained during MASLD progression and suggest that targeting NNMT may represent a promising therapeutic strategy for limiting hepatic inflammation and fibrosis.

## Materials and Methods

### Cell culture and treatments of HuH-7 cells and primary human hepatocytes

The human hepatoma cell line HuH-7 and cryopreserved primary human hepatocytes (PHHs) were used in this study. HuH-7 cells were cultured in Dulbecco’s Modified Eagle Medium (DMEM; Gibco) supplemented with 10% heat-inactivated fetal bovine serum (FBS; Gibco) and 100 U/mL penicillin-streptomycin (Gibco). Cells were maintained at 37°C in a humidified atmosphere containing 5% CO₂.

Cryopreserved PHHs were obtained from the Liver Cell Laboratory at the Division of Transplantation Surgery, Karolinska University Hospital, Sweden. Donor characteristics are summarized in **Table S1**. Cryopreserved PHHs were rapidly thawed in Cryopreserved Hepatocyte Recovery Medium (CHRM; Gibco) according to the manufacturer’s instructions and seeded onto collagen I-coated 6-well plates (Gibco). Cells were cultured in 500 mL Williams’ E Medium (Sigma-Aldrich) supplemented with 5 mL 200 mM L-glutamine, 10 mL 1 M HEPES buffer (Lonza), 10 μL 600 μM insulin (Sigma-Aldrich), 0.5 mL 50 mg/mL gentamicin (Sigma-Aldrich), 100 μL 250 μg/mL amphotericin B (Thermo Fisher Scientific), 50 μL 1 mM dexamethasone (Sigma-Aldrich) and 10% heat-inactivated FBS (Gibco).

For cytokine stimulation experiments, cells were treated with recombinant human TNFα (10 ng/mL), IL-1β (10 ng/mL), or IL-6 (10 ng/mL) for 6 h. Untreated cells served as controls. Both HuH-7 and PHHs were seeded at a density of 3×10^5^ cells per well in 6-well plates. All experiments were performed using biological replicates, unless otherwise stated. Detailed information regarding cytokines is provided in **Table S2**.

### LPS-treated mice

Male C57BL/6J mice aged 14-16 weeks were maintained on a standard chow diet. Mice were randomly assigned to experimental groups (n=3 per group) and administered lipopolysaccharide (LPS) intraperitoneally at a dose of 1 mg/kg body weight. Four hours following LPS injection, mice were sacrificed, and liver tissues were harvested for downstream analyses.

### siRNA-mediated knockdown in HuH-7 cells

HuH-7 cells were seeded at a density of 5×10⁴ cells per well in 24-well plates and allowed to adhere overnight at 37°C in a humidified atmosphere containing 5% CO₂. The following day, cells were transfected with control or gene-specific siRNA at a final concentration of 10 nM using Lipofectamine RNAiMAX (Thermo Fisher Scientific) according to the manufacturer’s instructions. After 6 h, the transfection medium was replaced with complete growth medium consisting of DMEM (Gibco) and 10% heat-inactivated FBS (Gibco). Cells were incubated for an additional 24 h before treatment with cytokines for 6 h. Subsequently, cells were harvested for downstream gene expression analyses. The sequences of all siRNAs used in this study are provided in **Table S3**.

### Nicotinamide (NAM) treatments in HuH-7 cells

For nicotinamide (NAM) (N0636-100G; Sigma-Aldrich) supplementation experiments, HuH-7 cells were treated with NAM at a final concentration of 500 μM. Following siRNA-mediated knockdown, cells were exposed to NAM in serum- and antibiotic-free DMEM either alone or in combination with cytokine treatment. After completion of the treatment period, cells were harvested for downstream gene expression analyses.

### Lentiviral-mediated NNMT overexpression in HuH-7 cells

HuH-7 cells were seeded at a density of 1×10⁴ cells per well in 96-well plates in DMEM (Gibco) supplemented with 10% FBS (Gibco) and allowed to adhere overnight at 37°C in a humidified atmosphere containing 5% CO₂. The following day, cells were transduced with either a control EGFP-expressing lentiviral vector or a human NNMT-overexpressing lentiviral vector (pLV[Exp]-EGFP/Puro-EF1A>hNNMT; VectorBuilder, Vector ID: VB900173-7714vmw) at a multiplicity of infection (MOI) of 4. Based on the viral titer (1×10⁶ PFU/mL), 10 μL of viral suspension was added per well. Cells were maintained under standard culture conditions, and transduction efficiency was monitored by fluorescence microscopy based on EGFP expression. Initial EGFP expression was observed 48 h post-transduction. To increase the proportion of transduced cells, cultures were maintained for approximately one week with regular medium replacement every 2-3 days and passaged as required. During this period, puromycin (4 μg/mL) (Gibco) was added to the culture medium to enrich for successfully transduced cells. Following expansion and selection, cells were exposed to the appropriate experimental conditions and harvested for downstream gene expression analyses.

### Human hepatic spheroid culture and treatments

Cryopreserved primary human liver cells were obtained from BioIVT (USA) and LifeNet Health (USA) and cultured as previously described by Youhanna et al. (29). PHHs and non-parenchymal cells (NPCs) were co-seeded at a PHH:NPC ratio of 6:1 into ultra-low attachment (ULA) 96-well plates (Corning) at a total density of 1,500 cells per well. Cells were cultured in Williams’ E medium (11 mM glucose) supplemented with 100 nM dexamethasone, 2 mM L-glutamine, 5.5 µg/mL transferrin, 6.7 ng/mL sodium selenite, 100 U/mL penicillin, 100 µg/mL streptomycin, and 10% FBS. Insulin was added at a final concentration of 10 ng/mL under lean conditions and 10 µg/mL under MASH conditions. Following spheroid formation, FBS was gradually withdrawn from the culture medium.

For the induction of steatosis/MASH, spheroids were cultured in medium supplemented with albumin-conjugated free fatty acids consisting of 240 µM palmitate and 240 µM oleate. Culture medium was refreshed every 2-3 days.

Intracellular lipid accumulation was quantified using the AdipoRed assay (Lonza, PT-7009) according to the manufacturer’s instructions. Briefly, spheroids were dissociated using 2.5% trypsin for 30 min at 37°C. Subsequently, 3.5 µL of AdipoRed reagent was added to each sample and mixed thoroughly. Fluorescence was measured at excitation and emission wavelengths of 485 nm and 572 nm, respectively. Lipid content was normalized to the corresponding control group and expressed as a percentage of control values.

Procollagen Iα1 levels in culture supernatants were determined using the Human Procollagen I alpha 1 DuoSet ELISA kit (R&D Systems, DY6220-05) following the manufacturer’s protocol.

PHHs were transfected with either NNMT-targeting siRNA or non-targeting control siRNA at a final concentration of 50 nM using Lipofectamine RNAiMAX (Invitrogen). siRNA-RNAiMAX complexes were prepared in serum-free Opti-MEM and incubated at room temperature for 20 min prior to use. Cells were subsequently combined with the transfection complexes and seeded into ULA 96-well plates for spheroid formation and further culture.

### Western blotting

HuH-7 cells were harvested and washed twice with phosphate-buffered saline (PBS) prior to lysis. Total protein was extracted using Pierce RIPA Buffer (Thermo Scientific), and protein concentrations were determined using the bicinchoninic acid (BCA) protein assay kit (Merck, 71285-M) according to the manufacturer’s instructions. Equal amounts of protein were separated by SDS-PAGE gels (Invitrogen) and transferred onto PVDF membranes (Cytiva, GE10600069). Membranes were blocked in StartingBlock T20 (PBS) Blocking Buffer (Thermo Scientific) for 1 h at room temperature and subsequently incubated with primary antibodies overnight at 4°C. Following washing, membranes were incubated with either peroxidase conjugated goat anti-mouse IgG secondary antibody (AP124P, Merck) or peroxidase conjugated donkey anti-rabbit IgG secondary antibody (NA934, Cytiva) for 1 h at room temperature. Protein bands were visualized using the ECL Prime Western Blotting Detection Reagent (Cytiva, RPN2232). A complete list of primary and secondary antibodies is provided in **Table S4**.

### SAM/SAH ELISA

Intracellular S-adenosylmethionine (SAM) and S-adenosylhomocysteine (SAH) levels were measured using a SAM/SAH ELISA Combo Kit (MyBioSource, Inc., San Diego, CA, USA; Cat. No. MBS169604) according to the manufacturer’s instructions. SAM and SAH concentrations were determined from standard curves, and the SAM/SAH ratio was calculated as a measure of cellular methylation potential.

### Quantitative PCR (qPCR) analysis

Total RNA was extracted from snap-frozen mouse liver tissues and human liver cells using the RNeasy Mini Kit (Qiagen, 74104) according to the manufacturer’s instructions. Complementary DNA (cDNA) was synthesized from total RNA using M-MLV Reverse Transcriptase (Life Technologies, 28025-021). qPCR was performed using the ABI Prism 7500 Real-Time PCR System (Applied Biosystems). Relative gene expression levels were calculated using the comparative cycle threshold (2^-ΔΔCt) method. For normalization, acidic ribosomal phosphoprotein P0 (Rplp0) was used as the housekeeping gene for mouse liver samples, whereas glyceraldehyde-3-phosphate dehydrogenase (GAPDH) was used as the housekeeping gene for human liver cells. Primer sequences used in this study are listed in **Table S5**.

### RNA sequencing (RNA-seq)

Total RNA was extracted from human liver cells as described above. RNA quality and integrity were assessed using the 4200 TapeStation system (Agilent). mRNA library preparation was performed using the NGS RNA Library Prep Set (PT042, Novogene) according to the manufacturer’s instructions. Libraries were sequenced as 150-bp paired-end reads (PE150) on the NovaSeq X Plus platform at Novogene (Cambridge, United Kingdom).

Raw sequencing reads were aligned to the human (GRCh38/hg38) reference genome using HISAT2. Gene-level read counts were generated using featureCounts v1.5.0-p3. Differential gene expression analysis was performed in R using the DESeq2 package. RNA-seq differential expression significant genes were defined as adjusted p-value < 0.05 and |log2 fold change 0.5|, unless otherwise stated. Additional downstream analyses and visualizations were performed using R.

In addition, processed gene count matrices from publicly available RNA-seq datasets GSE224012 and GSE135251 were downloaded from the Gene Expression Omnibus (GEO) database and incorporated into downstream analyses using R.

### Chromatin immunoprecipitation sequencing (ChIP-seq)

HuH-7 cells were crosslinked with 1% formaldehyde in phosphate-buffered saline (PBS) for 10 min at room temperature. Crosslinking was quenched by the addition of glycine to a final concentration of 0.125 M for 5 min. Cells were washed with ice-cold PBS supplemented with protease inhibitors and collected for chromatin preparation. Nuclei were isolated using sequential lysis buffers, and chromatin was fragmented by sonication using a Bioruptor Pico (Diagenode) to generate DNA fragments of approximately 200-500 bp. Chromatin immunoprecipitation was performed using 2 μg of anti-H3K27ac antibody (ab4729, Abcam) per sample, and immune complexes were captured using Protein A Dynabeads (Invitrogen) pre-incubated with the antibody overnight at 4°C. Following immunoprecipitation, beads were washed and crosslinking was reversed overnight at 65°C. ChIP DNA was purified using the ChIP DNA Clean and Concentrator Capped Zymo-Spin I purification kit (Zymo Research). For library preparation, ChIP DNA was processed using the ThruPLEX DNA-seq Kit (Takara Bio) according to the manufacturer’s instructions. Libraries were sequenced as 150-bp paired-end reads on the NovaSeq X Plus platform (Novogene; Cambridge, United Kingdom).

### ChIP-seq data analysis

ChIP-seq data analysis was performed using computational resources provided by the National Academic Infrastructure for Supercomputing in Sweden (NAISS) through project sens2024555 at UPPMAX. Raw sequencing reads (FASTQ files) generated by Novogene (Cambridge, United Kingdom) were aligned to the human reference genome (GRCh38/hg38) using Bowtie2. Peak calling was performed using the HOMER software package to identify regions enriched for H3K27ac.

For visualization in the Integrative Genomics Viewer (IGV 2.13.1), peak intensities were normalized to the total number of uniquely mapped reads and displayed as tags per 10 million mapped reads. Differential enrichment between experimental groups was determined using HOMER. Peaks with FDR < 0.05 and log2 fold change 0.5 were considered significantly differential. Genomic overlaps between peak sets were identified using BEDTools, and peak annotations were generated using HOMER. Signal intensity profiles and heatmaps were generated using deepTools. Subsequent analyses and visualizations were carried out in R.

### Integration of RNA-seq and ChIP-seq analyses

To integrate transcriptomic and epigenomic datasets, we performed an integrative transcription factor analysis based on the strategy described by Madsen et al. (26). Differential gene expression identified by RNA-seq was integrated with transcription factor motif enrichment analysis of differential H3K27ac regions performed using HOMER. Transcription factors supported by both differential gene expression and motif enrichment were prioritized as candidate regulators of the observed biological responses.

### Network visualization

Protein-protein interaction networks were generated using STRING and visualized in Cytoscape (v3.10.4). Fibrosis-associated gene modules were constructed from the selected candidate genes and used for downstream visualization and interpretation.

### Cleavage Under Targets and Tagmentation (CUT&Tag)

CUT&Tag experiments were performed following the protocol developed by the Henikoff laboratory (39). Briefly, cells were immobilized on Concanavalin A-coated magnetic beads (Polysciences) and incubated with antibodies against H3K4me2 (Abcam, ab7766), H3K27me3 (Millipore, 07-449), or H3K27ac (Abcam, ab4729), followed by incubation with the CUTANA Anti-Rabbit Secondary Antibody and pAG-Tn5 transposase (EpiCypher). DNA was purified, quantified using a Qubit fluorometer (Thermo Fisher Scientific), and sequencing libraries were prepared by PCR amplification using DNA/RNA UD Indexes (Illumina). Libraries were purified using SPRI beads (Beckman Coulter), quality controlled, and subjected to paired-end sequencing (150 bp) on the NovaSeq X Plus platform (Novogene, Cambridge, United Kingdom). A complete list of antibodies used in this study is provided in **Supplementary Table S4**.

### CUT&Tag data analysis

Sequencing data generated by Novogene (Cambridge, United Kingdom) were processed using standard bioinformatic pipelines. Raw sequencing reads (FASTQ files) were aligned to the human reference genome (GRCh38/hg38) using Bowtie2. Enriched chromatin regions were identified using SEACR, and RPKM-normalized coverage tracks were generated for visualization in the IGV 2.13.1. Differential enrichment analyses were performed in R using the DESeq2 or edgeR packages, depending on the dataset and analytical approach. Regions with adjusted p-value or FDR < 0.05 and log2 fold change 0.5 were considered significantly differential, unless otherwise stated. Downstream analyses and data visualization were performed in R.

### ATAC-seq (Assay for Transposase-Accessible Chromatin using sequencing) performance and analysis

ATAC-seq was performed according to the published protocol (25). Sequencing data generated by Novogene (Cambridge, United Kingdom) were processed using a standard bioinformatic pipeline. Raw sequencing reads were aligned to the human reference genome (GRCh38/hg38) using Bowtie2, and duplicate reads were removed prior to downstream analyses. Peaks were identified using HOMER, and RPKM-normalized coverage tracks were generated for visualization in IGV 2.13.1. Differential chromatin accessibility was assessed in R using the edgeR package. Regions with FDR < 0.05 and an absolute log2 fold change > 0.5 were considered differentially accessible. Downstream analyses and figures were performed in R.

## Statistical analysis

Statistical analyses were performed using GraphPad Prism 10 (GraphPad Software). Data are presented as mean ± standard deviation (SD). Biological replicates were used for each experiment. Comparisons between two groups were performed using two-tailed unpaired Student’s t-tests, whereas comparisons among multiple groups were analyzed using one-way analysis of variance (ANOVA) followed by appropriate post hoc tests, as indicated in the figure legends. All statistical tests were two-tailed, and p-value < 0.05 was considered statistically significant.

## Ethics

All animal experiments were approved by the Swedish Board of Agriculture under ethical permit number 05517-2022. The experiments were conducted in accordance with the International Guiding Principles for Biomedical Research Involving Animals developed by the Council for International Organizations of Medical Sciences (CIOMS). Mice were bred and maintained at the Center for Comparative Medicine at Karolinska Institutet and Karolinska University Hospital (PKL, Huddinge, Sweden).

The use of human liver tissue for primary hepatocyte isolation was approved by the Regional Ethical Review Board in Stockholm (Dnr: 2017/269-31). Human liver tissue was obtained from the Liver Cell Laboratory at the Division of Transplantation Surgery, Karolinska University Hospital, Sweden, in accordance with the approved ethical guidelines.

## Acknowledgments

Rongrong Fan was supported by grants from the EFSD Novo Nordisk future leaders award, the Swedish Research Council (2023–02311), the Swedish Cancer Society (232891 Pj), and the Karolinska Institutet Strategic Research Programme in Diabetes (SRP) Rolf Luft Grant. Eckardt Treuter was supported by grants from the Swedish Research Council (2022–00545), the Swedish Cancer Society (211582 Pj), and the Novo Nordisk Foundation (NNF22OC0078222). Lin Chen and Qi Li were supported by studentship from the Chinese Scholarship Council (CSC). Sviatlana Sukhanava, Philipp Valina Allo and Yue Zhu were supported by the Karolinska Institutet’s doctoral funding (KID). Volker Lauschke acknowledges support by the ERC Consolidator Grant 3DMASH (101170408), the Swedish Research Council (2021-02801, 2023-03015 and 2024-03401), Cancerfonden (24-3735Pj) and the Robert Bosch Foundation, Stuttgart, Germany.

## Author Contributions

R.R.F: Conceptualization, Methodology, Resources, Writing-Original Draft, Supervision, Project administration, Funding acquisition E.T: Conceptualization, Resources, Writing - Review & Editing, Supervision, Project administration, Funding acquisition S.S: Conceptualization, Methodology, Validation, Formal analysis, Investigation, Data curation, Writing-Original Draft, Visualization P.V.A: Methodology, Validation, Formal analysis, Investigation, Data curation, Visualization Y.Y.H., L.C: Methodology, Validation Y.Z: Formal analysis, Visualization S.Y: Methodology, Formal analysis, Visualization O.G.I: Methodology, Formal analysis Q.L: Formal analysis, Visualization E.E: Resources, Supervision V.M.L: Methodology, Resources, Supervision

## Declaration of interests

V.M.L is CEO and shareholder of HepaPredict AB, as well as co-founder and shareholder of Shanghai Hepo Biotechnology Ltd. The other authors declare no conflict of

## Data Availability

The sequencing data generated in this study have been deposited in the NCBI Gene Expression Omnibus (GEO) under accession numbers GSE345710, GSE341195, GSE341196, GSE341198, GSE341206, GSE339587, GSE341481.

## Notes

### Competing Interest Statement

VML is CEO and shareholder of HepaPredict AB, as well as co-founder and shareholder of Shanghai Hepo Biotechnology Ltd. The other authors declare no conflict of interest.

### Summary of Updates

We have repeated several key experiments, including Western blot analyses of histone methylation marks and CUT&Tag profiling, to further validate our findings. The new results indicate that, although NNMT deletion induces epigenetic alterations in liver cells, the predominant changes occur at H3K27ac rather than histone methylation marks. Furthermore, our new data suggest that these epigenetic changes are more closely associated with alterations in cellular NAD+ levels. Based on these additional experiments and the refined mechanistic insights, we have substantially revised the manuscript to ensure that its conclusions accurately reflect the experimental evidence.

## References

1. G. Targher, L. Valenti, C. D. Byrne, Metabolic Dysfunction–Associated Steatotic Liver Disease. New England Journal of Medicine 393, 683–698 (2025).

2. L. Hammerich, F. Tacke, Hepatic inflammatory responses in liver fibrosis. Nat Rev Gastroenterol Hepatol 20, 633–646 (2023).

3. S. Schuster, D. Cabrera, M. Arrese, A. E. Feldstein, Triggering and resolution of inflammation in NASH. Nat Rev Gastroenterol Hepatol 15, 349–364 (2018).

4. E. Kostallari, R. F. Schwabe, A. Guillot, Inflammation and immunity in liver homeostasis and disease: a nexus of hepatocytes, nonparenchymal cells and immune cells. Cell Mol Immunol 22, 1205–1225 (2025).

5. A.-M. Mirea, C. J. Tack, T. Chavakis, L. A. B. Joosten, E. J. M. Toonen, IL-1 Family Cytokine Pathways Underlying NAFLD: Towards New Treatment Strategies. Trends in Molecular Medicine 24, 458–471 (2018).

6. J. Schwärzler, P. Menghini, C. Dinarello, F. Cominelli, H. Tilg, IL-1 family of cytokines in gastrointestinal and liver disorders. Nat Rev Gastroenterol Hepatol 23, 29–43 (2026).

7. C. Garbers, S. Heink, T. Korn, S. Rose-John, Interleukin-6: designing specific therapeutics for a complex cytokine. Nat Rev Drug Discov 17, 395–412 (2018).

8. D. Schmidt-Arras, S. Rose-John, IL-6 pathway in the liver: From physiopathology to therapy. Journal of Hepatology 64, 1403–1415 (2016).

9. I. D. Vachliotis, S. A. Polyzos, The Role of Tumor Necrosis Factor-Alpha in the Pathogenesis and Treatment of Nonalcoholic Fatty Liver Disease. Curr Obes Rep 12, 191–206 (2023).

10. I. Goldstein, V. Paakinaho, S. Baek, M.-H. Sung, G. L. Hager, Synergistic gene expression during the acute phase response is characterized by transcription factor assisted loading. Nat Commun 8, 1849 (2017).

11. Y. Shen, et al., Decreased Hepatocyte Autophagy Leads to Synergistic IL-1β and TNF Mouse Liver Injury and Inflammation. 10.1002/hep.31209.

12. C.-Y. Zhang, et al., Nicotinamide N-methyltransferase in non-alcoholic fatty liver disease: Mechanistic insights and emerging therapeutic strategies. Archives of Biochemistry and Biophysics 772, 110558 (2025).

13. M. Komatsu, et al., NNMT activation can contribute to the development of fatty liver disease by modulating the NAD + metabolism. Sci Rep 8, 8637 (2018).

14. O. A. Ulanovskaya, A. M. Zuhl, B. F. Cravatt, NNMT promotes epigenetic remodeling in cancer by creating a metabolic methylation sink. Nat Chem Biol 9, 300–306 (2013).

15. C. Zhang, F. Macchi, E. Magnani, K. C. Sadler, Chromatin states shaped by an epigenetic code confer regenerative potential to the mouse liver. Nat Commun 12, 4110 (2021).

16. X. Lu, et al., NAD+ Metabolism Reprogramming Drives SIRT1-Dependent Deacetylation Inducing PD-L1 Nuclear Localization in Cervical Cancer. Advanced Science 12, 2412109 (2025).

17. Z. A. Wang, et al., Structural and enzymatic plasticity of SIRT6 deacylase activity. Journal of Biological Chemistry 301, 108446 (2025).

18. Q. Song, et al., ER stress-induced upregulation of NNMT contributes to alcohol-related fatty liver development. Journal of Hepatology 73, 783–793 (2020).

19. Y. Zhang, Z. Lu, W. Zeng, J. Zhao, X. Zhou, Two sides of NNMT in alcoholic and non-alcoholic fatty liver development. Journal of Hepatology 74, 1250–1253 (2021).

20. J. J. Babula, D. Bui, H. L. Stevenson, S. J. Watowich, H. Neelakantan, Nicotinamide N-methyltransferase inhibition mitigates obesity-related metabolic dysfunction. *Diabetes*, Obesity and Metabolism 26, 5272–5282 (2024).

21. O. Govaere, et al., Transcriptomic profiling across the nonalcoholic fatty liver disease spectrum reveals gene signatures for steatohepatitis and fibrosis. Sci Transl Med 12, eaba4448 (2020).

22. N. T. Doncheva, J. H. Morris, J. Gorodkin, L. J. Jensen, Cytoscape StringApp: Network Analysis and Visualization of Proteomics Data. J Proteome Res 18, 623–632 (2019).

23. Y. Feng, et al., ANGPTL8 deficiency attenuates lipopolysaccharide-induced liver injury by improving lipid metabolic dysregulation. Journal of Lipid Research 65, 100595 (2024).

24. L. Chen, et al., Sex-specific KDM6A-HNF4A-CREBH network controls lipoprotein cholesterol metabolism and atherosclerosis via epigenetic reprograming of hepatocytes. Nat Commun 17, 3945 (2026).

25. F. C. Grandi, H. Modi, L. Kampman, M. R. Corces, Chromatin accessibility profiling by ATAC-seq. Nat Protoc 17, 1518–1552 (2022).

26. J. G. S. Madsen, et al., Integrated analysis of motif activity and gene expression changes of transcription factors. Genome Res 28, 243–255 (2018).

27. G. Liang, et al., Nicotinamide N-methyltransferase and liver diseases. Genes Dis 10, 1883–1893 (2023).

28. M. P. Creyghton, et al., Histone H3K27ac separates active from poised enhancers and predicts developmental state. Proceedings of the National Academy of Sciences 107, 21931–21936 (2010).

29. S. Youhanna, et al., Chemogenomic Screening in a Patient-Derived 3D Fatty Liver Disease Model Reveals the CHRM1-TRPM8 Axis as a Novel Module for Targeted Intervention. Adv Sci (Weinh) 12, e2407572 (2025).

30. V. Arzumanian, M. Pyatnitskiy, E. Poverennaya, Comparative Transcriptomic Analysis of Three Common Liver Cell Lines. Int J Mol Sci 24, 8791 (2023).

31. D. E. Johnson, R. A. O’Keefe, J. R. Grandis, Targeting the IL-6/JAK/STAT3 signalling axis in cancer. Nat Rev Clin Oncol 15, 234–248 (2018).

32. B. Razani, et al., Negative Feedback in Noncanonical NF-κB Signaling Modulates NIK Stability Through IKKα-Mediated Phosphorylation. Sci Signal 3, ra41 (2010).

33. N. Oliva-Vilarnau, et al., Wnt/β-catenin and NFκB signaling synergize to trigger growth factor-free regeneration of adult primary human hepatocytes. Hepatology 79, 1337–1351 (2024).

34. K. Klöditz, E. Tewolde, Å. Nordling, M. Ingelman-Sundberg, Mechanistic, Functional, and Clinical Aspects of Pro-inflammatory Cytokine Mediated Regulation of ADME Gene Expression in 3D Human Liver Spheroids. Clin Pharmacol Ther 114, 673–685 (2023).

35. D. Kraus, et al., Nicotinamide N-methyltransferase knockdown protects against diet-induced obesity. Nature 508, 258–262 (2014).

36. M. Bockwoldt, et al., Identification of evolutionary and kinetic drivers of NAD-dependent signaling. Proc Natl Acad Sci U S A 116, 15957–15966 (2019).

37. S. Hong, et al., Nicotinamide N-methyltransferase regulates hepatic nutrient metabolism through Sirt1 protein stabilization. Nat Med 21, 887–894 (2015).

38. T. Bouras, et al., SIRT1 deacetylation and repression of p300 involves lysine residues 1020/1024 within the cell cycle regulatory domain 1. J Biol Chem 280, 10264–10276 (2005).

39. H. S Kaya-Okur, S. Henikoff, Bench top CUT&Tag v2. [Preprint] (2019). Available at: https://www.protocols.io/view/bench-top-cut-amp-tag-z6hf9b6 [Accessed 13 July 2026].

