## Supplementary figures for "Nicotinamide N-methyltransferase couples inflammation and epigenetic remodeling to hepatic fibrosis"

### Supplement Figures

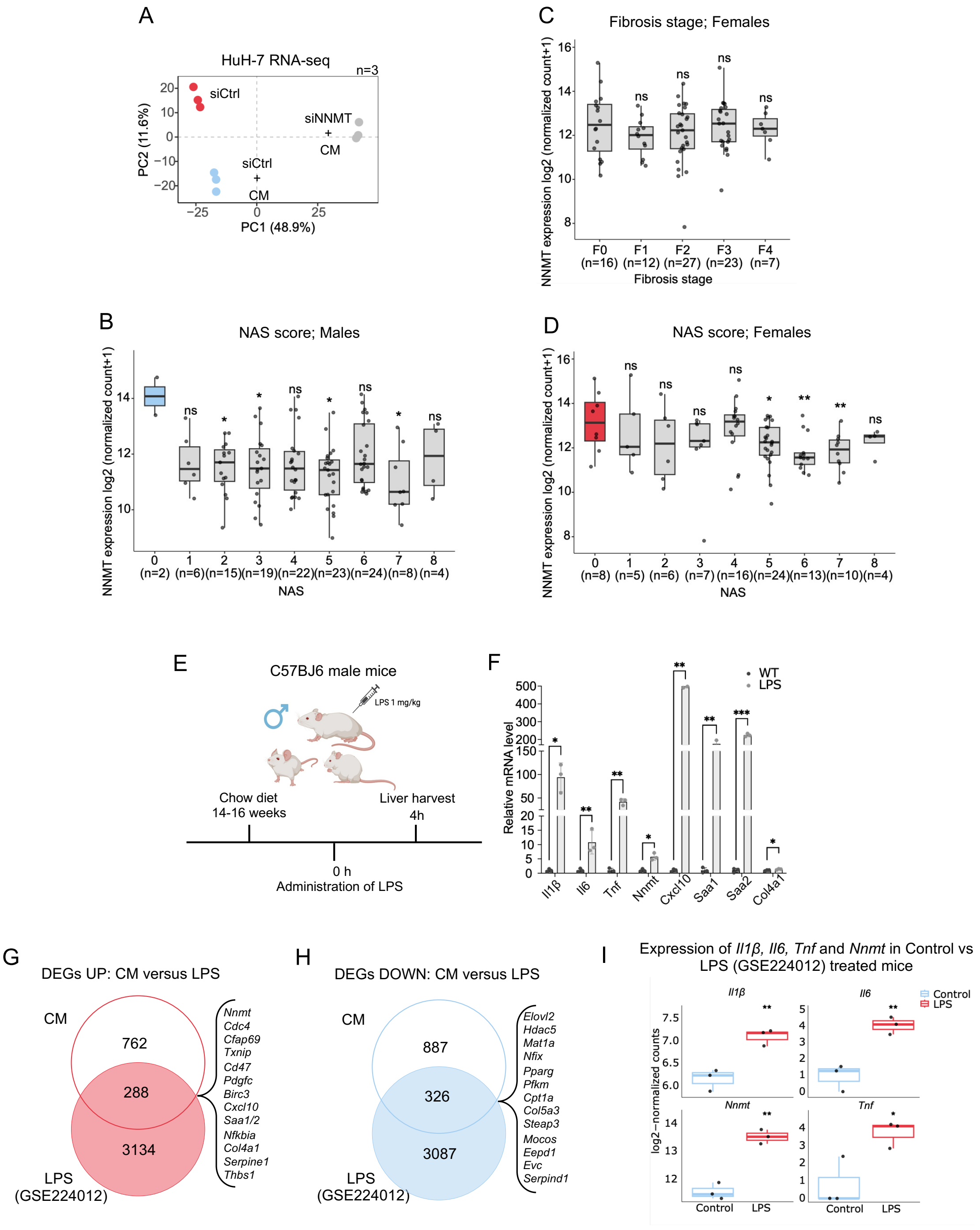
A
HuH-7 RNA-seq
n=3
siCtrl
siNNMT + CM
siCtrl + CM
PC1 (48.9%)
PC2 (11.6%)
B
NAS score; Males
NNMT expression log2 (normalized count+1)
0 (n=2) 1 (n=6) 2 (n=15) 3 (n=19) 4 (n=22) 5 (n=23) 6 (n=24) 7 (n=8) 8 (n=4)
NAS
C
Fibrosis stage; Females
F0 (n=16) F1 (n=12) F2 (n=27) F3 (n=23) F4 (n=7)
Fibrosis stage
D
NAS score; Females
0 (n=8) 1 (n=5) 2 (n=6) 3 (n=7) 4 (n=16) 5 (n=24) 6 (n=13) 7 (n=10) 8 (n=4)
NAS
E
C57BJ6 male mice
LPS 1 mg/kg
Chow diet
14-16 weeks
Liver harvest
4h
0 h
Administration of LPS
F
WT
LPS
Relative mRNA level
Il1β Il6 Tnf Nnmt Cxcl10 Saa1 Saa2 Col4a1
G
DEGs UP: CM versus LPS
CM
762
288
3134
LPS (GSE224012)
Nnmt Cdc4 Cfap69 Txnip Cd47 Pdgfc Birc3 Cxcl10 Saa1/2 Nfkbia Col4a1 Serpine1 Thbs1
H
DEGs DOWN: CM versus LPS
CM
887
326
3087
LPS (GSE224012)
Elovl2 Hdac5 Mat1a Nfix Pparg Pfkm Cpt1a Col5a3 Steap3 Mocos Eepd1 Evc Serpind1
I
Expression of Il1β, Il6, Tnf and Nnmt in Control vs LPS (GSE224012) treated mice
Il1β
Il6
Nnmt
Tnf
Control
LPS
log2−normalized counts

**Figure S1. *NNMT* expression in the GSE135251 patient cohort and conservation of *NNMT*-associated inflammatory responses in LPS-treated mouse liver, related to Figure 1.** (A) Principal component analysis of RNA-seq data from HuH-7 cells transfected with siCtrl or siNNMT under control or CM conditions (n = 3 per condition). (B) *NNMT* expression across NAS scores 0-8 in male patients of the GSE135251 cohort; each point represents one patient and the number of patients per group is indicated on the axis. (C) *NNMT* expression across fibrosis stages F0-F4 in female patients of the GSE135251 cohort. (D) *NNMT* expression across NAS scores 0-8 in female patients of the GSE135251 cohort. (E) Schematic of acute LPS treatment (1 mg/kg) in male C57BL/6J mice fed a chow diet for 14-16 weeks, with liver harvest 4 h after administration. Created with BioRender. (F) qPCR analysis of inflammatory and fibrosis-associated genes in liver tissue from control (WT) and LPS-treated mice; data are mean ± SD. (G, H) Overlap between genes upregulated (G) or downregulated (H) in CM-treated HuH-7 cells and in LPS-treated mouse liver from the public GSE224012 dataset; selected shared genes are listed. (I) Expression of *Il1β*, *Il6*, *Tnf* and *Nnmt* in WT and LPS-treated mouse liver from GSE224012. Comparisons in (B-D) are relative to the lowest score or stage. qPCR significance was determined by unpaired two-tailed t-test. *p < 0.05, **p < 0.01, ***p < 0.001; ns, not significant.

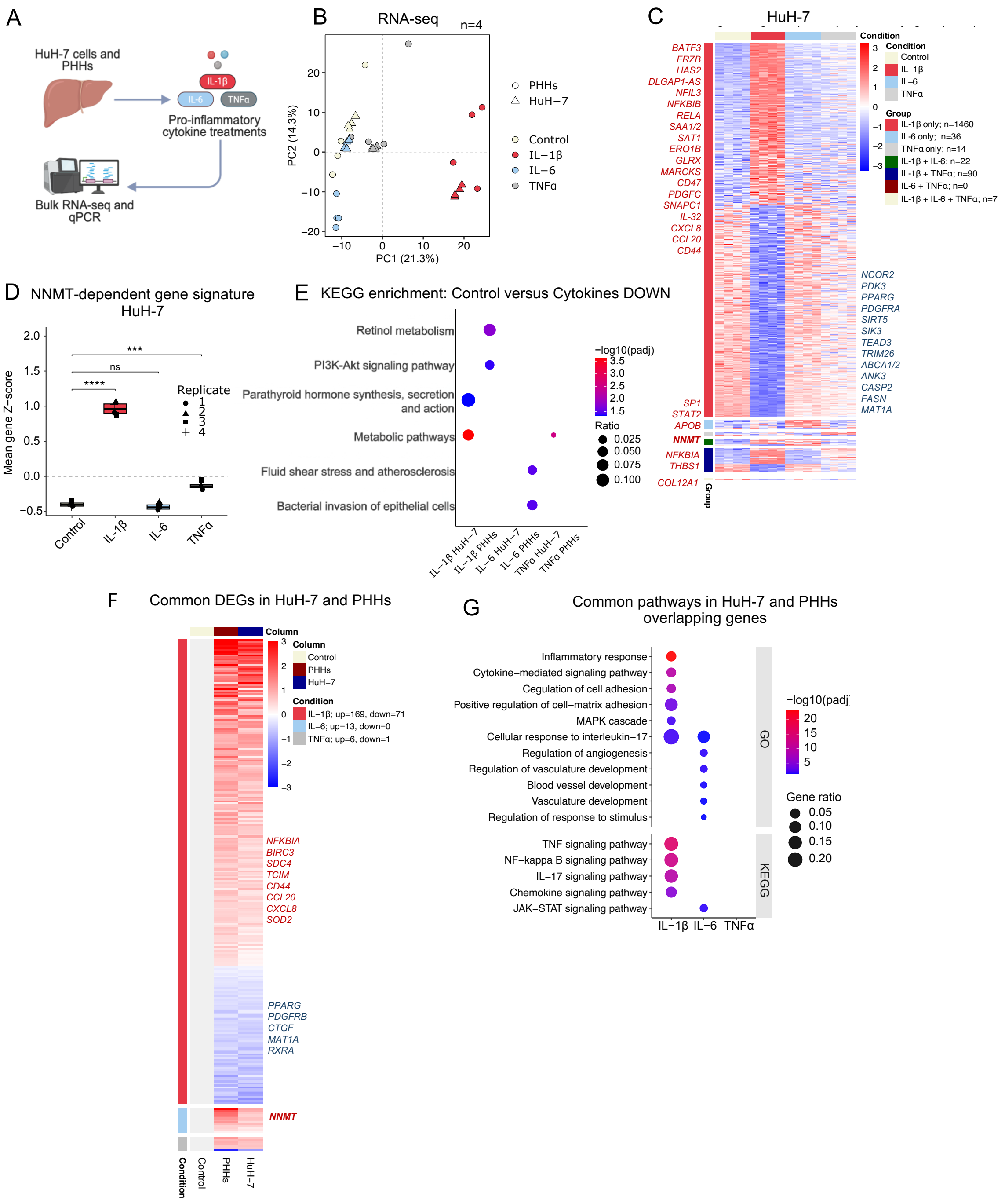

A
HuH-7 cells and PHHs
IL-1β
IL-6
TNFα
Pro-inflammatory cytokine treatments
Bulk RNA-seq and qPCR
B
RNA-seq
n=4
PC1 (21.3%)
PC2 (14.3%)
PHHs
HuH−7
Control
IL−1β
IL−6
TNFα
C
HuH-7
Condition
Control
IL−1β
IL−6
TNFα
Group
IL-1β only; n=1460
IL-6 only; n=36
TNFα only; n=14
IL-1β + IL-6; n=22
IL-1β + TNFα; n=90
IL-6 + TNFα; n=0
IL-1β + IL-6 + TNFα; n=7
BATF3
FRZB
HAS2
DLGAP1-AS
NFIL3
NFKBIB
RELA
SAA1/2
SAT1
ERO1B
GLRX
MARCKS
CD47
PDGFC
SNAPC1
IL-32
CXCL8
CCL20
CD44
SP1
STAT2
APOB
NNMT
NFKBIA
THBS1
COL12A1
NCOR2
PDK3
PPARG
PDGFRA
SIRT5
SIK3
TEAD3
TRIM26
ABCA1/2
ANK3
CASP2
FASN
MAT1A
D
NNMT-dependent gene signature
HuH-7
Mean gene Z−score
Replicate
***
ns
****
Control
IL-1β
IL-6
TNFα
E
KEGG enrichment: Control versus Cytokines DOWN
Retinol metabolism
PI3K-Akt signaling pathway
Parathyroid hormone synthesis, secretion and action
Metabolic pathways
Fluid shear stress and atherosclerosis
Bacterial invasion of epithelial cells
−log10(padj)
Ratio
IL−1β HuH−7
IL−1β PHHs
IL−6 HuH−7
IL−6 PHHs
TNFα HuH−7
TNFα PHHs
F
Common DEGs in HuH-7 and PHHs
Column
Control
PHHs
HuH−7
Condition
IL−1β; up=169, down=71
IL−6; up=13, down=0
TNFα; up=6, down=1
NFKBIA
BIRC3
SDC4
TCIM
CD44
CCL20
CXCL8
SOD2
PPARG
PDGFRB
CTGF
MAT1A
RXRA
NNMT
G
Common pathways in HuH-7 and PHHs overlapping genes
Inflammatory response
Cytokine−mediated signaling pathway
Cegulation of cell adhesion
Positive regulation of cell−matrix adhesion
MAPK cascade
Cellular response to interleukin−17
Regulation of angiogenesis
Regulation of vasculature development
Blood vessel development
Vasculature development
Regulation of response to stimulus
TNF signaling pathway
NF−kappa B signaling pathway
IL−17 signaling pathway
Chemokine signaling pathway
JAK−STAT signaling pathway
GO
KEGG
−log10(padj
Gene ratio
IL−1β
IL−6
TNFα

**Figure S2. Cytokine-specific transcriptional responses in HuH-7 cells and primary human hepatocytes, related to Figure 2.** (A) Experimental overview of HuH-7 cells and PHHs treated with IL-1β, IL-6 or TNFα followed by bulk RNA-seq and qPCR analyses. Created with BioRender. (B) Principal component analysis of RNA-seq data from HuH-7 cells and PHHs following cytokine treatment (n = 4 per condition). (C) Heatmap of differentially expressed genes in HuH-7 cells following cytokine treatment relative to control. Genes are grouped by cytokine-specific and shared responses, with the number of genes per group given in the key; colours indicate row-scaled expression Z-scores. (D) *NNMT*-dependent gene signature score in cytokine-treated HuH-7 cells, calculated as the mean row-scaled Z-score across the signature genes; each point represents one biological replicate and comparisons are relative to control. (E) KEGG enrichment of genes downregulated by individual cytokines in HuH-7 cells and PHHs; dot size indicates gene ratio and colour indicates -log10 adjusted p-value. (F) Heatmap of common cytokine-responsive genes shared between HuH-7 cells and PHHs, with the number of shared up- and downregulated genes per cytokine given in the key; colours indicate row-scaled expression Z-scores. (G) GO and KEGG enrichment of common cytokine-responsive genes shared between HuH-7 cells and PHHs; dot size indicates gene ratio and colour indicates -log10 adjusted p-value. The *NNMT*-dependent gene signature score in (D) was calculated as the mean Z-score across *NNMT*-dependent genes for each replicate, and statistical comparisons were performed using t-tests with Benjamini–Hochberg correction for multiple comparisons. ***$p < 0.001$, ****$p < 0.0001$; ns, not significant.

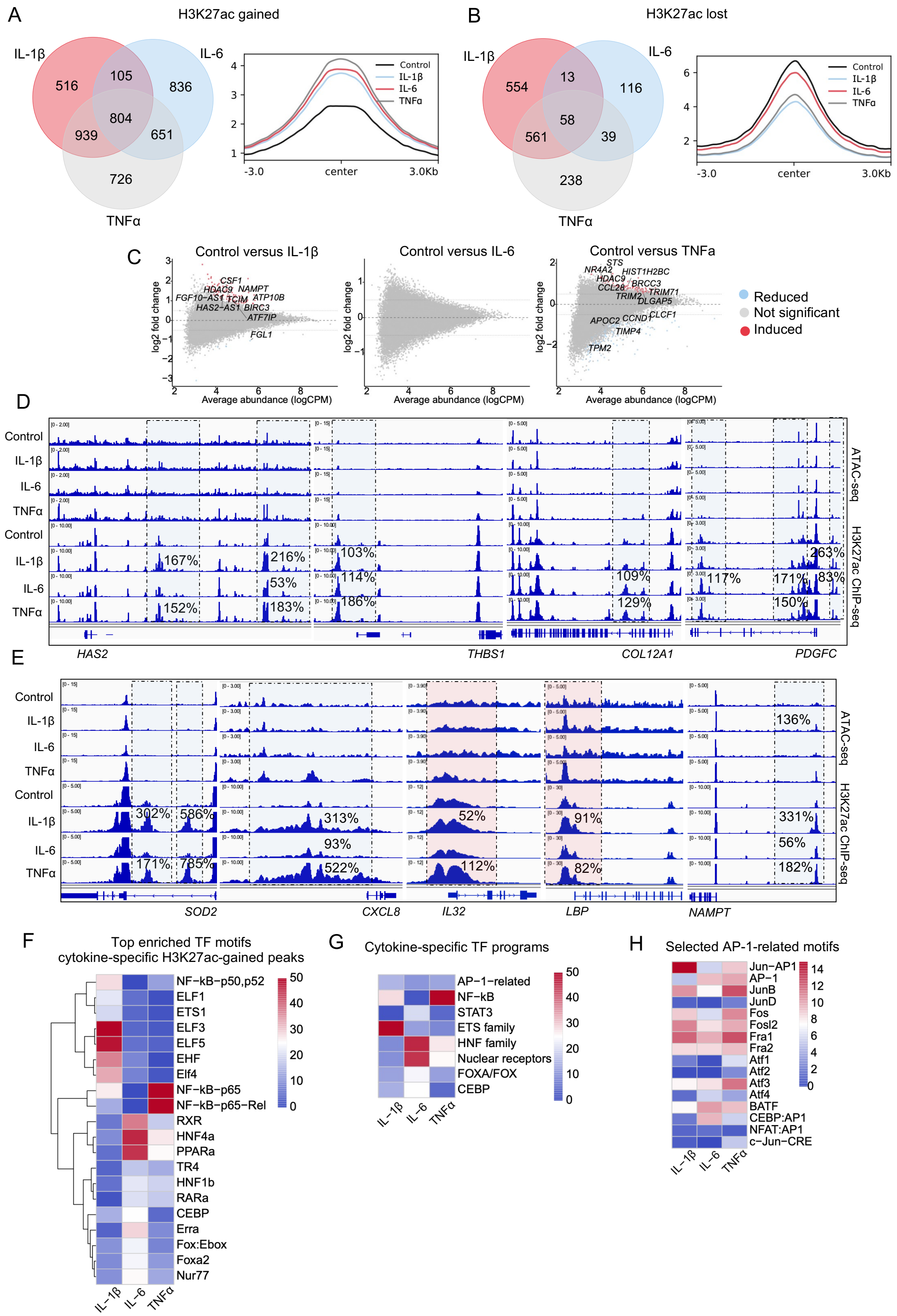
A
H3K27ac gained
IL-1β
IL-6
TNFα
516
105
836
804
939
651
726
Control
IL-1β
IL-6
TNFα
-3.0
center
3.0Kb
B
H3K27ac lost
554
13
116
58
561
39
238
C
Control versus IL-1β
Control versus IL-6
Control versus TNFa
log2 fold change
Average abundance (logCPM)
Reduced
Not significant
Induced
D
ATAC-seq
H3K27ac ChIP-seq
HAS2
THBS1
COL12A1
PDGFC
E
SOD2
CXCL8
IL32
LBP
NAMPT
F
Top enriched TF motifs
cytokine-specific H3K27ac-gained peaks
NF−kB−p50,p52
ELF1
ETS1
ELF3
ELF5
EHF
Elf4
NF−kB−p65
NF−kB−p65−Rel
RXR
HNF4a
PPARa
TR4
HNF1b
RARa
CEBP
Erra
Fox:Ebox
Foxa2
Nur77
G
Cytokine-specific TF programs
AP−1−related
NF−kB
STAT3
ETS family
HNF family
Nuclear receptors
FOXA/FOX
CEBP
H
Selected AP-1-related motifs
Jun−AP1
AP−1
JunB
JunD
Fos
Fosl2
Fra1
Fra2
Atf1
Atf2
Atf3
Atf4
BATF
CEBP:AP1
NFAT:AP1
c−Jun−CRE

**Figure S3. Cytokine stimulation remodels H3K27ac at inflammatory and fibrosis-associated regulatory regions, related to Figure 3.** (A, B) Venn diagrams showing H3K27ac-gained (A) and H3K27ac-lost (B) peaks following IL-1β, IL-6 or TNFα treatment, with H3K27ac signal profiles shown for the corresponding regions. (C) MA plots of differential chromatin accessibility from ATAC-seq in IL-1β-, IL-6- and TNFα-treated HuH-7 cells compared with control cells; red, increased accessibility; blue, decreased accessibility; grey, not significant. (D, E) Representative genome browser tracks showing H3K27ac ChIP-seq and ATAC-seq signal at selected fibrosis-associated (D) and inflammatory/metabolic (E) loci; red shading indicates promoter regions and blue shading indicates intronic or intergenic regulatory regions. Percentages indicate the descriptive change in signal relative to control. (F-H) Motif enrichment heatmaps of cytokine-specific H3K27ac-gained peaks showing top enriched transcription factor motifs (F), selected cytokine-specific transcription factor programs (G) and selected AP-1-related motifs (H); colours indicate absolute log p-values.

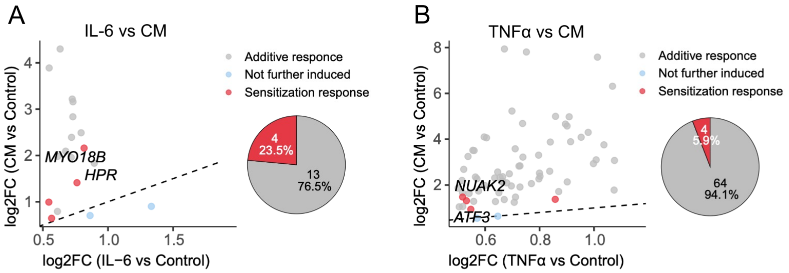


**Figure S4. Limited sensitization of IL-6- and TNFα-responsive genes by cytokine mix treatment in HuH-7 cells, related to Figure 4.** (A, B) RNA-seq comparison of IL-6-responsive (A) or TNFα-responsive (B) genes following single-cytokine stimulation versus CM treatment in HuH-7 cells. Genes are classified as additive responses, not further induced, or sensitization responses. Pie charts summarise the proportion of sensitized genes among overlapping cytokine-responsive genes.

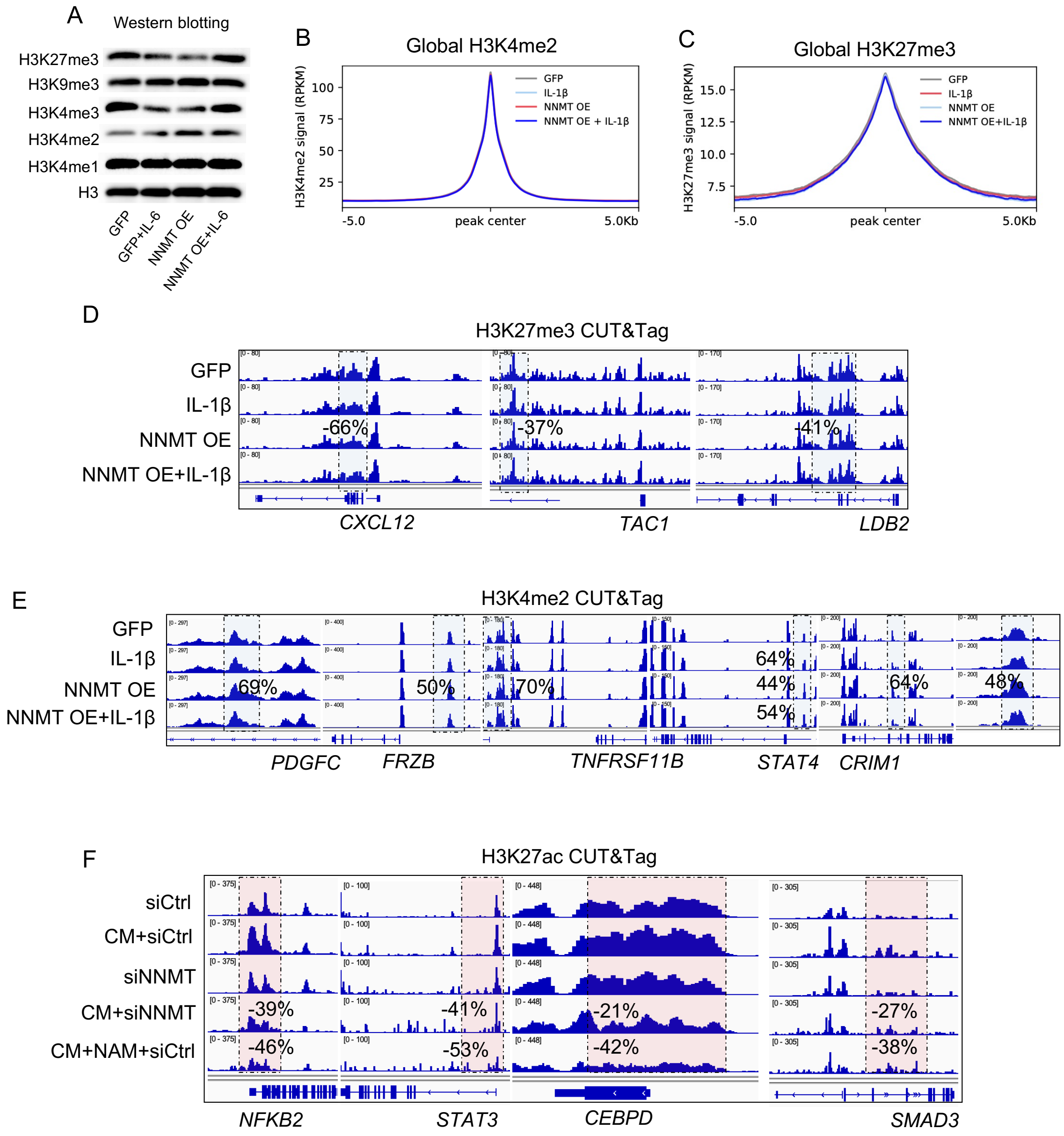


**Figure S5. *NNMT* overexpression is associated with selective H3K4me2 remodeling and limited H3K27me3 changes, related to Figure 5.** (A) Representative western blots of H3K27me3, H3K9me3, H3K4me3, H3K4me2, H3K4me1 and total H3 in GFP, GFP+IL-1β, *NNMT* overexpression (OE) and *NNMT* OE+IL-1β HuH-7 cells. (B, C) Average global H3K4me2 (B) and H3K27me3 (C) CUT&Tag signal profiles across experimental conditions. (D) Representative H3K27me3 CUT&Tag genome-browser tracks at the *CXCL12*, *TAC1* and *LDB2* loci. (E) Representative H3K4me2 CUT&Tag genome-browser tracks at the *PDGFC*, *FRZB*, *TNFRSF11B*, *STAT4* and *CRIM1* loci. (F) Representative H3K27ac CUT&Tag genome-browser tracks at the *NFKB2*, *STAT3*, *CEBPD* and *SMAD3* loci. For H3K27ac CUT&Tag, biological replicate numbers are siCtrl, n = 3; CM+siCtrl, n = 3; siNNMT, n = 2; CM+siNNMT, n = 1; and CM+NAM+siCtrl, n = 1. Percentages indicate the change in H3K27ac signal in CM+siNNMT and CM+NAM+siCtrl relative to mean CM+siCtrl signal.

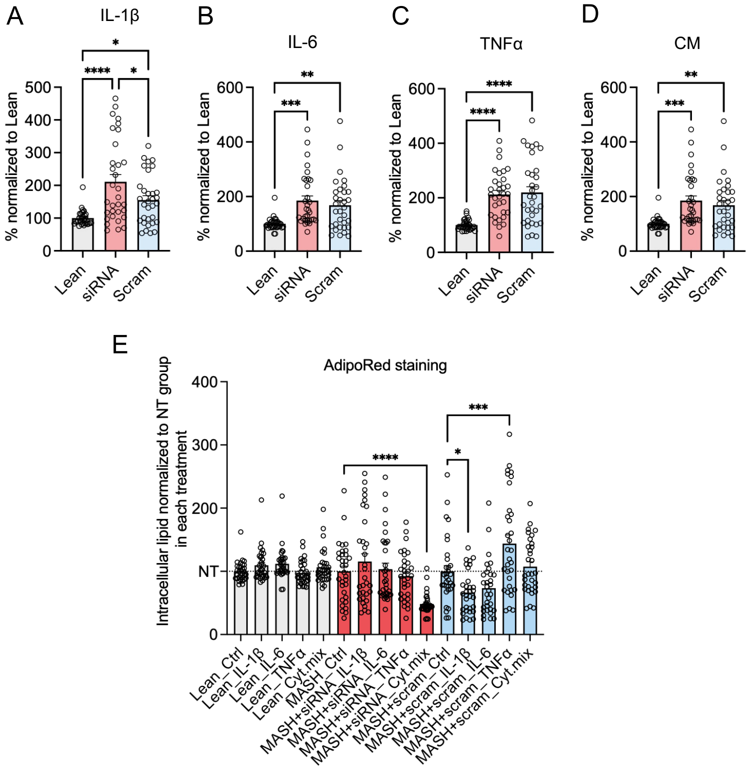


**Figure S6. *NNMT* knockdown selectively modulates lipid accumulation in cytokine-treated human liver spheroids, related to Figure 6.** (A-D) Quantification of intracellular lipid accumulation by AdipoRed assay in liver spheroids following IL-1β (A), IL-6 (B), TNFα (C) or CM (D) treatment under lean or MASH conditions with scrambled control siRNA (scram) or *NNMT* siRNA. Values are normalised to lean spheroids. (E) Intracellular lipid accumulation across treatment conditions measured by AdipoRed staining and normalised to untreated controls within each treatment group. Data are presented as mean ± SD. Statistical comparisons were performed using one-way ANOVA with multiple-comparison correction. *$p < 0.05$, **$p < 0.01$, ***$p < 0.001$, ****$p < 0.0001$.

**Tables**

**Table S1. Characteristics of primary human hepatocyte donors.**

| **Donor #** | **Tissue source** | **Sex** | **Age (years)** |
|---|---|---|---|
| 1 | Liver resection | Male | 75 |
| 2 | Liver resection | Male | Unknown |
| 3 | Donor | Male | 34 |
| 4 | Donor | Male | 41 |

**Table S2. Pro-inflammatory cytokines.**

| Reagent | Supplier | Catalog Number |
|---|---|---|
| IL-1β | Sigma-Aldrich | H6291-10UG |
| IL-6 | Sigma-Aldrich | SCU0001-10UG |
| TNFα | Sigma-Aldrich | H8916-10UG |

**Table S3. siRNA sequences.**

| Target | Sense (5'-3') | Antisense (5'-3') |
|---|---|---|
| siLuc | CGUACGCGGAAUACUUCGA(dT)(dT) | UCGAAGUAUUCCGCGUACG(dT)(dT) |
| siNNMT | GCCAGAUUCUUAAGCACCU(dT)(dT) | AGGUGCUUAAGAAUCUGGC(dT)(dT) |
| siJUN | GAUGGAAACGACCUUCUAU(dT)(dT) | AUAGAAGGUCGUUUCCAUC(dT)(dT) |

**Table S4. Antibodies for CUT&Tag, ChIP-seq and Western blot.**

| Antibody | Supplier | Catalog Number |
|---|---|---|
| β-actin | Abcam | ab5694 |
| H3K27me3 | Sigma-Aldrich | 07-449 |
| H3K9me3 | Abcam | Ab8898 |
| H3K4me3 | Abcam | ab8580 |
| H3K4me2 | Abcam | ab7766 |
| H3K4me1 | Abcam | ab8895 |
| H3K27ac | Abcam | ab4729 |
| H3 | Abcam | ab1791 |
| Normal Rabbit IgG | Cell Signaling Technology | 2729S |
| Rabbit IgG HRP Linked Whole Ab | Cytiva | GENA934 |
| Goat Anti-Mouse IgG Antibody, Peroxidase Conjugated, H+L | Sigma-Aldrich | AP124P |

**Table S5. qPCR primer sequences.**

| **Human primers** | **Forward (5'-3')** | **Reverse (5'-3')** |
|---|---|---|
| *JUN* | TCCAAGTGCCGAAAAAGGAAG | CGAGTTCTGAGCTTTCAAGGT |
| *NNMT* | TGGTGACCTATGTGTGTGATCT | CCCTGGCTTCAGTAGGCTG |
| *CCL20* | TGCTGTACCAAGAGTTTGCTC | CGCACACAGACAACTTTTTCTTT |
| *IL-32* | TGGCGGCTTATTATGAGGAGC | CTCGGCACCGTAATCCATCTC |
| *NFKBIA* | TGTGCTTCGAGTGACTGACC | TCACCCCACATCACTGAACG |
| *LBP* | ACTGGAGAGCAGGATTTGCG | AATGTCGGCGAAACTGTCAAT |
| *CXCL8* | TTTGCCAAGGAGTGCTAAAGA | AACCCTCTGCACCCAGTTTTC |
| *THBS1* | GCCATCCGCACTAACTACATT | TCCGTTGTGATAGCATAGGGG |
| *SERPINE* | ACCGCAACGTGGTTTTCTCA | TTGAATCCCATAGCTGCTTGAAT |
| *COL12A1* | AGCTGAGGCAGACATTGTGTT | CCTCCTTTGTACGGCAAGTTT |
| *HAS2* | AAGAACAACTTCCACGAAAAGGG | GGCTGGGTCAAGCATAGTGT |
| *SDC4* | GGACCTCCTAGAAGGCCGATA | AGGGCCGATCATGGAGTCTT |
| *SOD2* | GCTGGAAGCCATCAAACGTG | TGCTCCCACACATCAATCCC |
| *CPT1A* | ATCAATCGGACTCTGGAAACGG | TCAGGGAGTAGCGCATGGT |
| *NAMPT* | ATCCTGTTCCAGGCTATTCTGT | CCCCATATTTTCTCACACGCAT |
| *ELOVL7* | GCCTTCAGTGATCTTACATCGAG | AGGACATGAGGAGCCAATCTT |
| *IL-18* | CAGATCGCTTCCTCTCGCAA | CCAGGTTTTCATCATCTTCAGCTAT |
| *FBLN5* | CTCACTGTTACCATTCTGGCTC | GACTGGCGATCCAGGTCAAAG |
| *CSF1* | TGGCGAGCAGGAGTATCAC | AGGTCTCCATCTGACTGTCAAT |
| *IL-1RN* | TCCCCATGGCTTTAGAGACGA | GACAGGCACATCTTCCCTCC |
| *ACSL1* | CTTATGGGCTTCGGAGCTTTT | CAAGTAGTGCGGATCTTCGTG |
| *FCAMR* | GAGAAGTGGACTACTCCAGGC | GCGAAAGAAGAACCTTGTAGCAG |
| *GAPDH* | GTCTCCTCTGACTTCAACAGCG | ACCACCCTGTTGCTGTAGCCAA |

| **Mouse primers** | **Forward (5'-3')** | **Reverse (5'-3')** |
|---|---|---|
| *Il1β* | AAATACCTGTGGCCTTGGGC | CTTGGGATCCACACTCTCCAG |
| *Il6* | GCTGGAGTCACAGAAGGAGTGGC | TCTGACCACAGTGAGGAATGTCCA |
| *Tnf* | AAGCCTGTAGCCCACGTCGTA | GGCACCACTAGTTGGTTGTCTTTG |
| *Nnmt* | GCCCCACCATCTATCAGCTTC | ACGCCTCAACTTCTCCTCCT |
| *Cxcl10* | CCAAGTGCTGCCGTCATTTTC | GGCTCGCAGGGATGATTTCAA |
| *Saa1* | GCGAGCCTACACTGACATGA | TGTCTGTTGGCTTCCTGGTC |
| *Saa2* | TATGATGCTGCCCAAAGGGG | CATGGTGTCCTCGTGTCCTC |
| *Col4a1* | CCTGGCACAAAAGGGACGA | ACGTGGCCGAGAATTTCACC |
| *Rplp0* | ACTGGTCTAGGACCCGAGAAG | TCCCACCTTGTCTCCAGTCT |
